# Unraveling the Phylogenetic Complexities of *Ischyropsalis* C.L. Koch, 1839 (Opiliones: Dyspnoi: Ischyropsalididae) in the North Iberian Peninsula, with the description of three new species

**DOI:** 10.64898/2026.01.09.698449

**Authors:** Ricardo López-Alonso, Esteban Pacual-Parra, Lucía Labrada, Carlos G. Luque, Eduardo Cires, Andrés Arias, Claudia González-Toral

## Abstract

*Ischyropsalis* is a genus of harvestmen inhabiting both terrestrial and caves in central and southern Europe. The accepted number of species is controversial due to feature similarities and their strong sexual dimorphism. The northern part of the Iberian Peninsula harbours 11 accepted taxa, four of which are terrestrial, while the other seven exclusively live in caves. Some of these species have not been included in the only phylogenetic study conducted on this genus, so their taxonomic status has not been evaluated yet. In this context, we aim to unravel the phylogenetic relationships among the taxa within this genus, from the northern part of the Iberian Peninsula, by conducting morphological and phylogenetic analyses based on mitochondrial (COI) and nuclear (EF1-α) molecular markers. Phylogenetic analyses revealed three previously undescribed species from karst caves in Cantabria and Asturias, herein described as *I. damiani* sp. nov., *I. impressa* sp. nov. and *I. aguerana* sp. nov.

## Introduction

Opiliones, commonly known as harvestmen, are a monophyletic order of arachnids formed by four suborders, 70 families, around 1691 genera, and more than 6700 described species (summarized in Kury *et al*. 2023). Dyspnoi, one of the major clades, includes the monogeneric sub-family Ischyropsalidinae Simon, 1879 (Schönhofer 2013). Ischyropsalidinae is a monotypic and morphologically distinct west Palearctic subfamily, being its genus *Ischyropsalis* C.L. Koch, 1839 restricted to the central and southern European mountain regions (i.e. the Alps, Carpathians, Apennines and the northern part of the Iberian Peninsula) (Schönhofer *et al*. 2015). The members of this genus, characterised by large bodies and chelicerae and a distinct penis morphology, have been reported to be difficult to find due to their habitat preferences, which in some cases include caves (Luque & Labrada 2012; Schönhofer 2013; Schönhofer *et al*. 2015). Recent morphological and molecular studies revealed that the Basque-Cantabrian Mountains (northern Iberian Peninsula), the Alps and the Pyrenees are centres of endemism for this genus (Schönhofer 2013; Schönhofer *et al*. 2015). The genus *Ischyropsalis* is the only known representative of its subfamily, which is therefore monotypic. It was established with *Ischyropsalis kollari* C.L. Koch, 1839 as the type species by subsequent designation (Thorell 1876: 467). Its definition was later modified and expanded with the addition of further species. At present, the genus includes 22 valid species and was revised by Schönhofer (2013). At the time of his study, Martens (1969b) could not examine the few species known from the Cantabrian Mountains, which are very closely related and strictly allopatric. Consequently, parts of his treatment of this fauna remained hypothetical. These points were later corrected by the works of E. Dresco, C. Prieto, and C.G. Luque (see Luque & Labrada 2012; Schönhofer 2013, and references therein).

The taxonomy of this genus is based on a combination of the male cheliceral apophysis, including the glandular area and the penis morphology. (Martens 1969a, 1969b; Prieto 1990a; Luque 1991; Luque & Labrada 2012; Schönhofer *et al*. 2015). The number of species has varied during the decades due to morphological similarities, which could be convergent in the case of the genitalia according to Martens (1969b), and the strong sexual dimorphism displayed by the *Ischyropsalis* taxa (Martens 1978; Schmidt *et al*. 2024). Thus, several authors have described many species (e.g. C.F. Roewer in 1950 described 30 new species), some of which have been found to be synonymous species (e.g. Roewer 1950; Helversen & Martens 1972) (see Schönhofer 2013 for specifications). In this context, Martens (1969b) and Dresco (1970), the former focused mainly on alpine species and the latter on Iberian species, made some of the most significant contributions based on morphology to the taxonomy of this genus, although their evolutionary and biogeographic hypotheses seldom coincided (Schönhofer *et al*. 2015). In addition, the phylogenetic study of Schönhofer et al. (2015) focused on most of the 22 valid species recognised by Schönhofer (2013). Using mitochondrial and nuclear molecular markers (*Cytochrome Oxidase Subunit I* (COI), *Elongation Factor 1 Alpha* (EF1-α) and *28S ribosomal RNA* (28S rRNA)), the authors showed that the morphological characters employed in current systematics were not lineage-associated. This result provided further support for the observations of Martens (1969b). Schönhofer *et al*. (2015) recovered three major phylogenetic lineages within *Ischyropsalis*: 1) the basal monospecific *I. hellwigii* group (*I. hellwigii* subsp. *hellwigii* (Panzer, 1793) and *I. hellwigii* subsp. *lucantei* (Simon, 1879)); 2) the *I. manicata* group formed by species occurring in the western Alps, the Apennines and the Carpathians (*I. carli* Lessert, 1905*, I. alpinula* Martens, 1978, *I. manicata* Koch, 1869 and *I. adamii* Canestrini, 1873); 3) the “morphologically heterogenous” Alpine-Iberian group, formed by the Alpine clade (*I. dentipalpis* Dresco, 1959, *I. kollari* Koch, 1839, *I. lithoclasica* Martens & Schönhofer, 2010, *I. muellneri* Harmann, 1898, *I. ravasinii* Hadzi, 1942 and *I. strandi* Kratochvil, 1936), and 4) the Iberian clade (*I. luteipes* Simon, 1872, *I. pyrenaea* Simon, 1872, *I. dispar* Simon, 1872, *I. magdalenae* Simon, 1881, *I. navarrensis* Roewer, 1950*, I. nodifera* Simon, 1879 and *I. robusta* Simon, 1872). However, it should also be highlighted that, although no lineage-specific morphological features were identified by Schönhofer et al. (2015), the molecular analyses showed that the morphological traits traditionally used in systematics are not lineage-associated and may therefore be misleading (Martens 1969b; Prieto 1990a; Luque 1991; Luque & Labrada 2012; Schönhofer & Martens 2015). For example: the size, position and shape of the glandular fields, often associated with knob-like apophyses exhibited on the dorsal and/or distal side of the field, are species-specific features and serve as main characters for species delineation and identification (Martens 1969b; Schönhofer & Martens 2015). Although comprehensive, the study of Schönhofer et al. (2015) did not include in their molecular analysis the Iberian taxa *I. hispanica* Roewer, 1953, *I. petiginosa* Simon, 1913, *I. cantabrica* Luque & Labrada, 2012 and *I. gigantea* Dresco, 1968. Prieto *et al*. (2013) also studied the Iberian *Ischyropsalis* taxa using molecular methods based on COI. Nevertheless, this study corresponds to a conference poster therefore, the genetic and morphological data of this study are not publicly available. Furthermore, the number and names of the outgroups used to build the presented *COI* topology were not specified, limiting the comparison of our *COI* results with the topology presented by Prieto et al. (2013).

Schönhofer (2013) included 11 taxa occurring in the Iberian Peninsula. Five show a epigean behaviour (*I. lucantei*, *I. luteipes*, *I. nodifera*, *I. petiginosa* and *I. hispanica*), while six are hypogean (*I. pyrenaea*, *I. navarrensis*, *I. dispar*, *I. magdalenae*, *I. cantabrica* and *I. gigantea*) (Luque & Labrada 2012). The seven Iberian taxa molecularly studied by Schönhofer et al. (2015) (*I. luteipes*, *I. pyrenaea*, *I. dispar*, *I. magdalenae*, *I. navarrensis*, *I. nodifera* and *I. robusta*) were found in the Iberian clade, which comprised of two inner clades, the Pyrenean and Cantabrian clades, and whose members are characterised by extraordinarily long and slender chelicerae. Within this group, the differences in size and morphology of both chelicerae and the bristle field on the basal cheliceral article, located on the distomedial side, are used to differentiate between species (Schönhofer *et al*. 2015).

Some of the untreated Iberian taxa occur in the karst areas of the Cantabrian Mountain Range, located in northern Iberian Peninsula (Sendra, 2023). Many of these karstic areas are of special importance for the Cantabrian and Asturian biodiversity, as they host highly specialized and endemic cave-dwelling fauna, with some taxa presenting unique, unusual or even bizarre morphological, behavioural or ecological adaptations (Labrada et al., 2010, in press; Salgado *et al*. 2012; Angus *et al*. 2012; Luque & Labrada 2012, 2017; Faille *et al*. 2021; Sendra 2023; Fresneda *et al*. 2025). In these karstic area, two endemic troglobiont Ischyropsalis species occur: the Cantabrian species *I. gigantea* and *I. cantabrica* as well as the undescribed forms treated herein. Moreover, they are located within a latitudinal belt (ca. 42° to 46° N in Europe) a high biodiversity of cave-dwelling terrestrial fauna (‘mid-latitude biodiversity ridge’) resulting from historical, mostly climatic, conditions (Culver *et al*. 2006). On the other hand, the species status of some other Iberian taxa occurring in this area, such as the terrestrial *I. hispanica*, remains controversial as this was synonymized as *I. nodifera* by Martens (1969b), while Prieto (1990a) reassigned this taxon as valid species. In contrast, *I. petiginosa*, *I. gigantea* and *I. cantabrica*, though clearly distinguishable morphologically and grouped into the Cantabrian clade based on their distribution and morphological traits (Luque 1991; Luque & Labrada 2012), still lack molecular validation to confirm their phylogenetic independence (Schönhofer *et al*. 2015).

In this context where the phylogenetic position and taxonomic status of at least four classically recognised species of the Cantabrian group (*I. hispanica*, *I. petiginosa*, *I. nodifera* and *I. cantabrica*) has not been molecularly validated and three putative new species have been found in an area with high *Ischyropsalis* diversity, we aim to clarify the specific status of these taxa and to describe three new species found in different caves in the north of Iberian Peninsula using morphological and molecular methods.

## Materials and methods

### Study area

The Cantabrian Range has an irregular but extensive area (approximately 20,000 km^2^) of carbonate outcrops of Palaeozoic (pre-Carboniferous and Carboniferous) limestones and Mesozoic (Jurassic and Cretaceous) limestones and dolomites, with a large number of caves and therefore potential cavernicolous habitats (Sendra 2023). For instance, in the provinces of Cantabria (5,321 km^2^) and Asturias (10,604 km^2^) alone, accounting for 3.14% of the territory of Spain, approximately 20,000 caves have been documented (as visualized at the Spanish Caving Federation (FEE), and Cantabrian (FCE) and Asturian (FESPA) Caving Federations websites). Twenty-seven hydrogeological units have been identified by the Geological Survey of Spain in the Cantabrian region (García de la Noceda *et al*. 1999). In order to maximise the range of potential cavernicolous habitats, we have sampled three units that differed in terms of age of the rock formation, number of caves, altitude, distance from present sea-shore and geographic longitude. Fifteen caves were selected for this study in the Cantabrian region, divided into two areas: from East to West, selected hydrogeological units were Cantabria province (Figs. 1-2) and partly Somiedo-Trubia-Pravia in Asturias province (Fig. 3).

**Figure 1.**
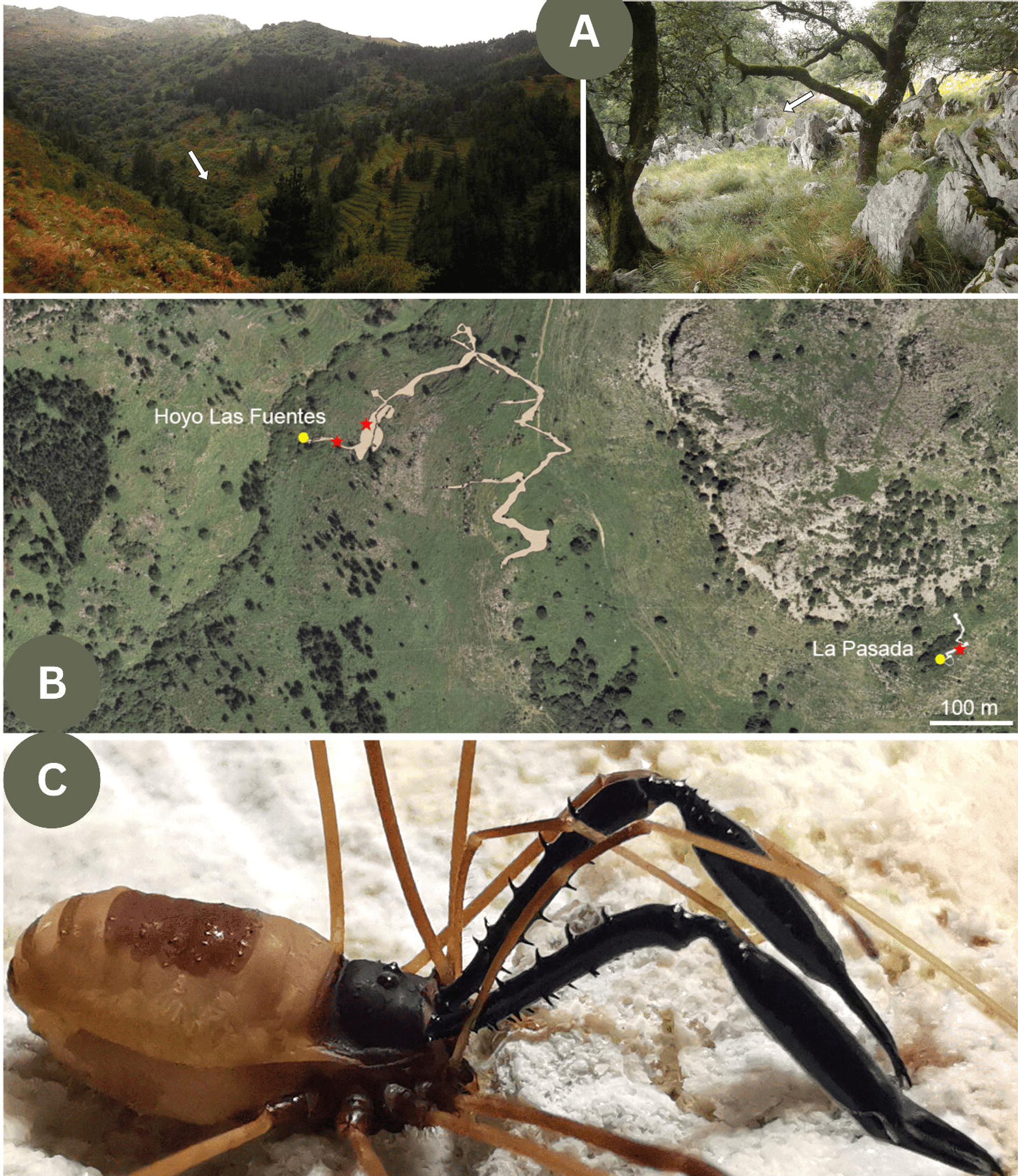
Geographic distribution of *Ischyropsalis impressa* sp. nov. (yellow dots) and *I. aguerana* sp. nov. (white dots) in northeastern Cantabrian Mountains. A) Location of study area and map of the Karst in Spain at 1:1,000,000-scale where karstified carbonate rocks emerge. B) LiDAR visualisation of the study area showing Cantabria–Biscay border (solid yellow line). The orange-brown area represents the range of limestone outcrops from the Urgonian formation (of Aptian age, Early Cretaceous) at 1:50,000-scale. Location of the caves surveyed: 1. La Baja; 2. Hoyo de Peñaflor; 3. Perelada; 4. Hoyo Molino; 5. Hoyo Menor-Tocinos; 6. Hoyo Menor-Rejuyo; 7. Hoyo de la Cubilla; 8. La Lastrilla; 9. Covacho del Cogorio; 10. Hoyo de las Rebolligas; 11. La Pasada; 12. Hoyo de las Fuentes; 13. La Tabla II; 14. Gueriza.

**Figure 2.**
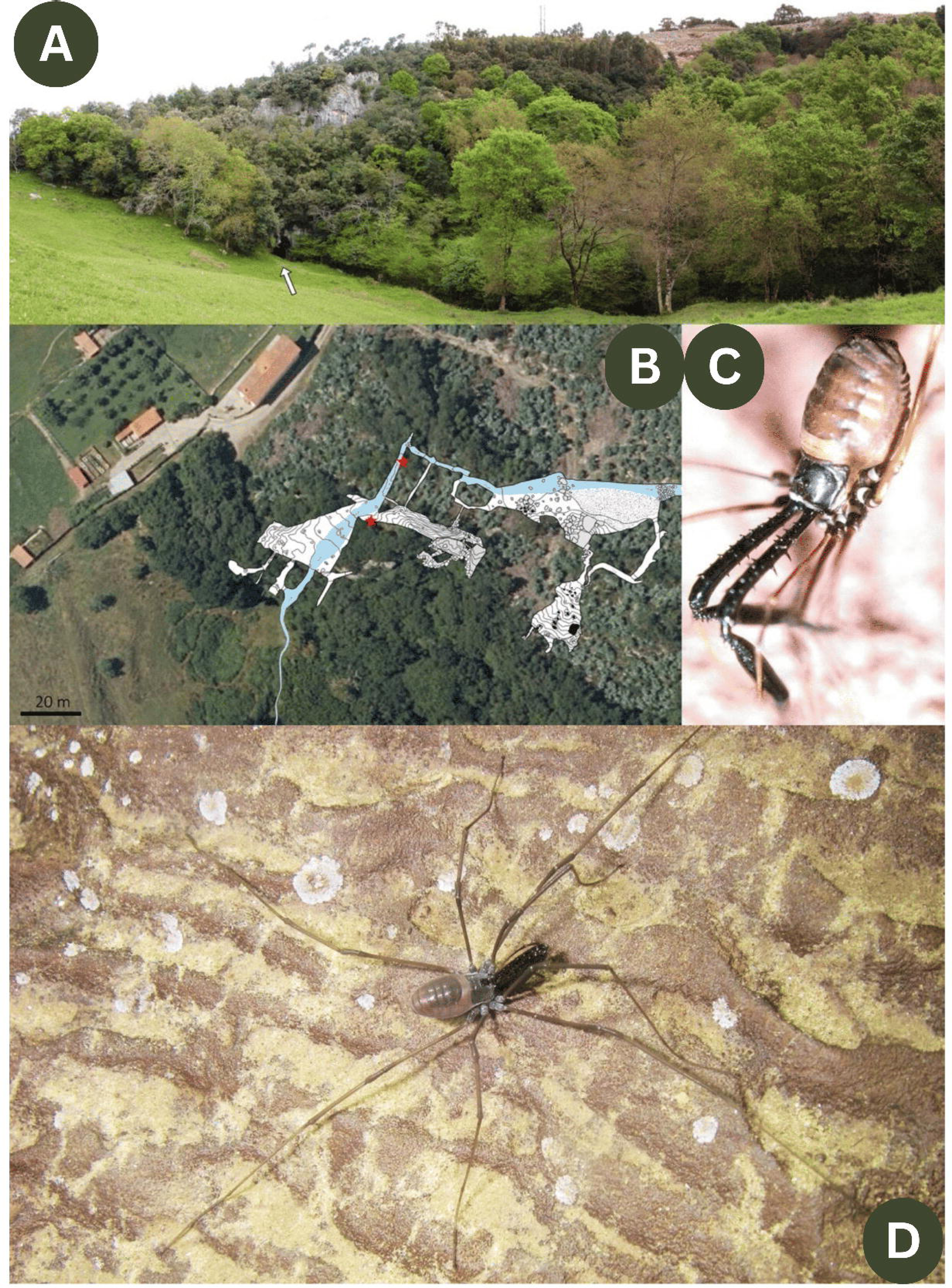
Distribution of the four troglobiont *Ischyropsalis* species in Cantabria and geological map at 1:50,000-scale the outcrops of the rock systems (in gray) which contain limestone (solid yellow line is Cantabria–Burgos–Biscay border): *cantabrica* (A), *impressa* sp. nov. (B), *aguerana* sp. nov. previously published as Ischyropsalis aff. dispar (Fig. 55 in Luque & Labrada, 2012) (C) and *gigantea* (D). Male cheliceral apophysis on distal end of basal article with basal part of second article of right chelicera (medial view) and distal part of penis (dorsal view) are shown.

**Figure 3.**
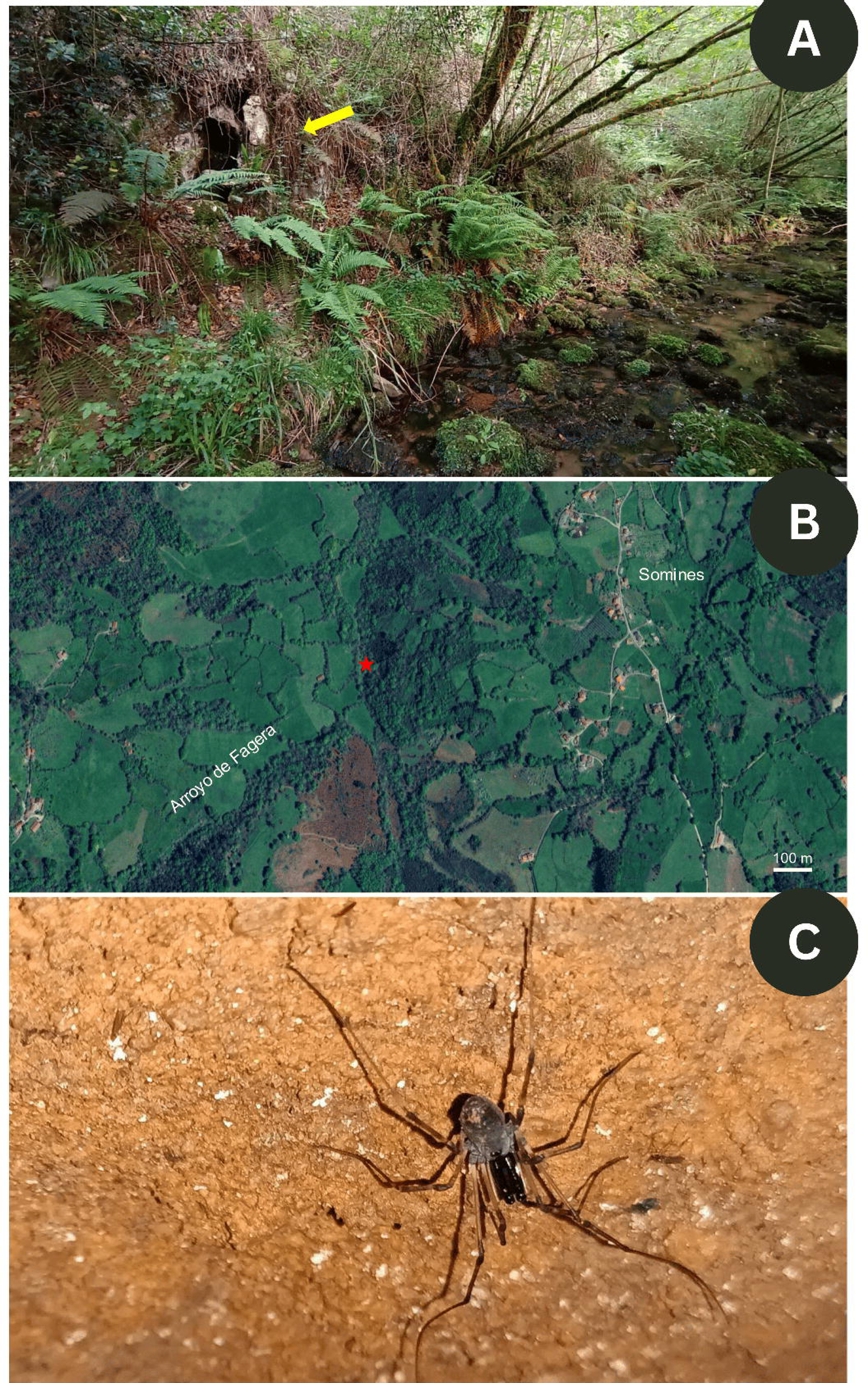
Type location of *Ischyropsalis damiani* sp. nov. in Asturias region (indicated by star). A) Location of study area and map of the Karst in Spain at 1:1,000,000-scale where karstified carbonate rocks emerge (karst areas distribution is based on Geological Survey of Spain; IGME, Spanish acronym). B) Location of Elvira Cave and areas of Oviedo with bedrock in which karst processes occur (the range of limestone outcrops of Carboniferous age is represented in grey, at a scale of 1:50,000)

### Specimens of investigation and specimens’ collection

The specimens were collected using pitfall traps and direct capture. They were preserved either in 75% ethanol (EtOH) or in absolute EtOH and deposited in the following institutions and collections: MNCN, National Museum of Natural Sciences, Madrid, Spain; CRBA (collection of M. Rambla), Animal Biodiversity Resource Centre at Barcelona University, Spain; ZUPV, Department of Zoology at Basque Country University, Spain; BOS (subcollection of Opiliones: BOS-OPI, Spanish acronym), Department of Organisms and Systems Biology at Oviedo University, Asturias, Spain; CJM, collection of J. Martens, Institute of Zoology, University of Mainz, Germany; CGL, private collection of C.G. Luque, Cantabria, Spain. Field collecting permits were issued to C.G. Luque and L. Labrada by Regional Government of Cantabria (Spain). The species used in the phylogenetic analyses were collected from the type localities of the species.

### SEM Ultrastructural Analysis

Specimens for Scanning Electron Microscope (SEM) were air-dried, mounted on aluminium stub, metalized with gold (16 nm of deposit) in an Argon atmosphere with bio-rad SC-515 by the “Sputter coating” method and with acceleration velocities of 15 kv. Initially, some of the specimens were dehydrated in an ethanol series reaching 100% to be dried by CO2 critical point in a pressure chamber. Stubs were examined and micrographs provided under high vacuum in a Philips XL-20 SEM at the National Museum of Natural Sciences (Spain) and in a JEOL JSM-6610LV SEM at the University of Oviedo. Chelicerae and genitalia of male and female specimens were excised, mounted on slides with clove oil, and illustrated using a Carl Zeiss IV binocular microscope with a *camera lucida*. Description and abbreviations format follows Luque & Labrada (2012). Morphological nomenclature followed Martens (1969b) for male internal genitalia and shape of the chelicera apophyses, like effectively used in previous works (Luque & Labrada 2012; Schönhofer 2013). The final versions of the drawings were prepared using Adobe Photoshop software version 7.0. All measurements are in millimetres (mm) with a precision of 0.01 mm and were taken employing a micrometer disc and by using the same optical devices. We give the average value followed by the minimum and maximum range in brackets.

### DNA extraction and amplification

DNA was extracted from the legs of the male specimens (including individuals from species type localities) following the instructions of the DNeasy Blood and Tissue kit (Qiagen, Hilden, Germany) as done in previously studies such as Schönhofer *et al*. (2015). The obtained DNA was stored at -20°C. The nuclear marker *Elongation Factor 1 Alpha* (EF1-α) was amplified using the pair of primers OP2BSAB and OPRC4 and the PCR amplification protocol described by Hedin *et al*. (2010). The *Cytochrome Oxidase Subunit I* (COI) mitochondrial marker was amplified by using the primers C1-J-1718SPIDERA and C1-N2776SPIDER and the PCR protocol described in Vink *et al*. (2005). For those samples with fragmented DNA, the internal set of primers of Leray *et al*. (2013), mlCOIintF and mlCOIintR, and the universal primers of Folmer *et al*. (1994), HCO2198 and LCO1490, were used, following the PCR conditions described by Leray *et al*. (2013). The PCR products were visualized in agarose gel. The ones with the expected size (see Table 1 for details) were sent to Macrogen (Macrogen, Madrid, Spain) for Sanger sequencing.

**Table 1.**
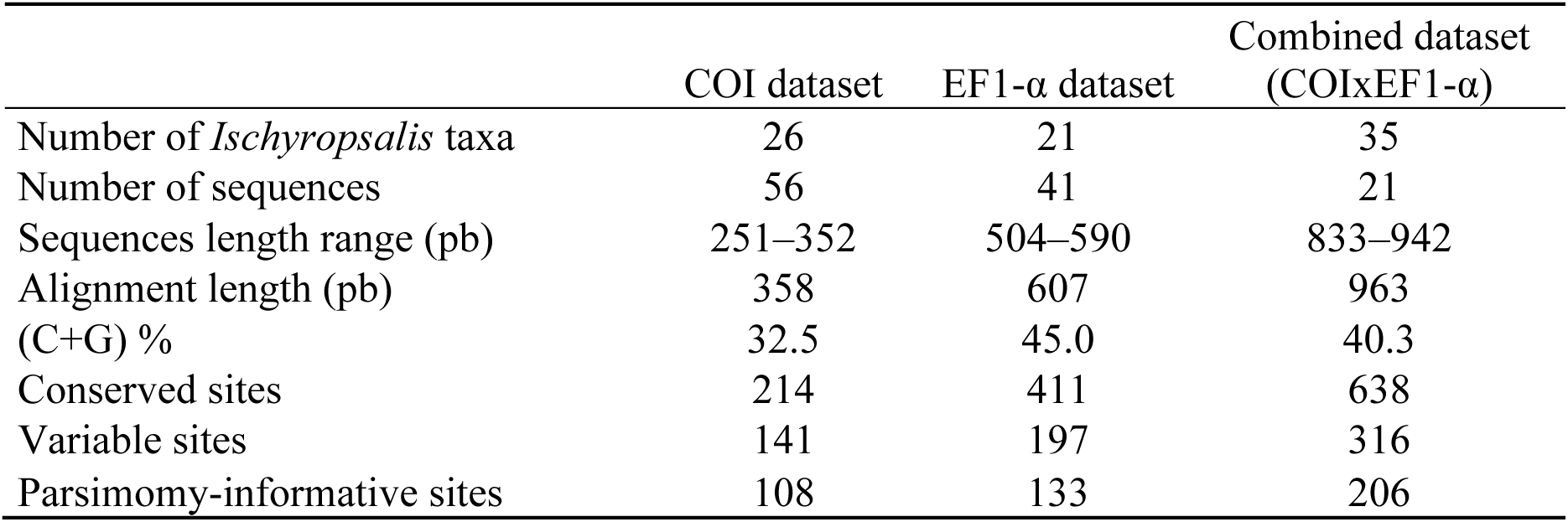
Main features of the mitochondrial *Cytochrome Oxidase Subunit I* (COI), *Elongation Factor 1 Alpha* (EF1-α) and Combined (COIxEF1-α) alignments used for the *Ischyropsalis* phylogenetic analyses. The alignment positions that contain at least two different nucleotides were considered as variables, while when each of the different nucleotides occurring in apposition had a minimum frequency of two, those positions were considered parsimonious-informative.

### Phylogenetic analyses

The obtained sequences were reviewed and edited using Geneious Prime V.2023.2.1 Software (Kearse *et al*. 2012). When the lower peaks were a third part of the total length of a higher peak and occurred in the reverse and forward sequences, double peaks were considered as ‘valid’.

The edited sequences were used to generate three different datasets, one for each molecular marker (the EF1-α dataset, the COI dataset, and both of them) together with the *Ischyropsalis* GenBank sequences used in previous phylogenetical analyses of this genus (Schönhofer *et al*. 2015) (see Supplementary Material 1). The Ischyropsalididae family outgroups were the same as in Schönhofer *et al*. (2015): *Acuclavella merickeli* Shear, 1986, *Acuclabella sheari* Richart & Hedin, 2013, *Acuclavella makah* Richart & Hedin, 2013 and *Ceratolasma trincantha* Goodnight & Goodnight, 1942. The alignments were performed by Multiple Sequence Comparison by Long-Expectation algorithm (MUSCLE) as implemented in Jalview 2.11.1.3 (Edgar 2004; Waterhouse *et al*. 2009), using the Akaike Information Criterion (AIC) implemented in jModelTest 2.1.10 (Akaike 1974; Darriba *et al*. 2012) to determine the best-fitting substitution. The General Time Reversible model with gamma distribution (GTR + G) (Tavaré 1986; Yang 1994) was the best-fitting model for the COI, EF1-α and both dataset markers. Additionally, separate EF1-α datasets were generated for exon-only sequences and for combined exon–intron sequences, (following Schönhofer et al., 2015 methodology) and independent phylogenetic analyses were performed for each dataset. The resulting topologies were compared with those obtained from the full dataset and are provided in Supplementary Material 3.

Two different methods were used to determine the phylogenetic relationships within *Ischyropsalis*: Maximum Likelihood (ML) as implemented in IQtree V. 1.6.12 webserver (Nguyen *et al*. 2015; Trifinopoulus *et al*. 2016) and Bayesian Inference (BI) as implemented in Mr. Bayes 3.2 (Ronquist *et al*. 2012). The ML trees were built by creating an initial Neighbour-Joining (NJ) tree and using the Nearest-Neighbour Interchange (NNI) method for branch rearranging (Robinson 1971; Moore *et al*. 1973). Branch support was estimated by 10,000 ultrafast bootstrap replications (BT) (Minh *et al*. 2013; Chernomor *et al*. 2016; Hoang *et al*. 2017). Two parallel runs of Markov Chain Monte Carlo (MCMC), consisting in 1 cold and 3 hot chains each, were used for the BI analyses, which ran for 10,000,000 generations and had a burning fraction of 0.25 (Hastings 1970). The Posterior Probability (PP) was used to estimate the statistical branch support in the BI analyses (Hastings 1970; Metropolis *et al*. 1953). Whenever the statistical branch support was below 50 %, tree branches were collapsed.

## Results

### Taxonomy

Class Arachnida Cuvier, 1812 Order Opiliones Sundevall, 1833

Suborder Dyspnoi Hansen & Sørensen, 1904 Family Ischyropsalididae Simon, 1879 Subfamily Ischyropsalidinae Simon, 1879 Type genus: *Ischyropsalis* C.L. Koch, 1839

Diagnosis (emended from Rozwałka et al. 2012 and Schönhofer 2013)

Defined by peculiarities of penis glans morphology, the enlarged chelicerae with glandular fields, located distally on the basal article in males of most species and supported by molecular phylogenetic evidence. Body egg-shaped, uniformly black or brownish-black and greyish or brownish (occasionally white). Mostly lacking dorsal armature, smooth dorsum, often glossy, mostly with scutum laminatum or parvum (rarely scutum intermedium); in few species opisthosomal areas raised to small, pointed bumps. Two or more spine-like metapeltidial structures, arranged in a transversal row. Ocularium without spine; eyes separated by distinct furrow. Chelicerae massive and enlarged, distinctly longer than the body (can be up to ca. one and half to twice body length). In most species, male chelicerae are spineless or have only short piliferous tubercles; female chelicerae possess spines only. Male chelicerae with dorsal or dorso-medial secretion-extruding field indicated by bristle-field in the distal end of basal article. In most species glandular fields of males associated with knob-like apophyses. Long and thin palps. Legs of medium to long length, robust and unarmed. Club-shaped penis with simple truncus, basal two-thirds portion with a single intrinsic muscle. Short muscular tendon, the base of truncus divided into two short root-like structures. Glans conical to inflated. The sclerotised ventral plate is short, from rhombic to bi-lobed at the base. Sclerotised area of glans with several defined regions exhibiting dense fields of very thick, long and curved backward-pointing setae. Glans gradually tapering into a long stylus, which is simple and pointed, bent at the base but without contortion or additional structures. Female with very short ovipositor, not segmented, with few setae in the apical area, *receptaculum seminis* with numerous (4–6, rarely 10) finger-like ampullae, basally surrounded with funnel-shaped structure.

### Species descriptions

*Ischyropsalis impressa* sp. nov. Luque & Labrada, 2025 ZooBank taxon LSID: F225B713-7333-4BFE-A09A-910EB3271261 Figs. 1–2, 4-6, 17G

### Diagnosis

A species of the westernmost *dispar*-group (see Luque & Labrada 2012) with short and slender chelicerae, which can be told apart by morphological characters and by allopatric range. Males are discernible by the chelicerae, which show a weak distal bulge on the basal article (Figs. 4D, 5B–5D), as well as a mace-shaped distal end with an extended bristle area (Figs. 5B, 7), and by their distinct penis morphology (Figs. 5H, 6B–6D).Females can be distinguished by the presence of a relatively short, or sometimes absent, dorsal spine at the distal end of the basal article, together with several large dorsal and ventral spines on the basal cheliceral article (Figs. 4C, 5E–5G, see arrows).

**Figure 4.**
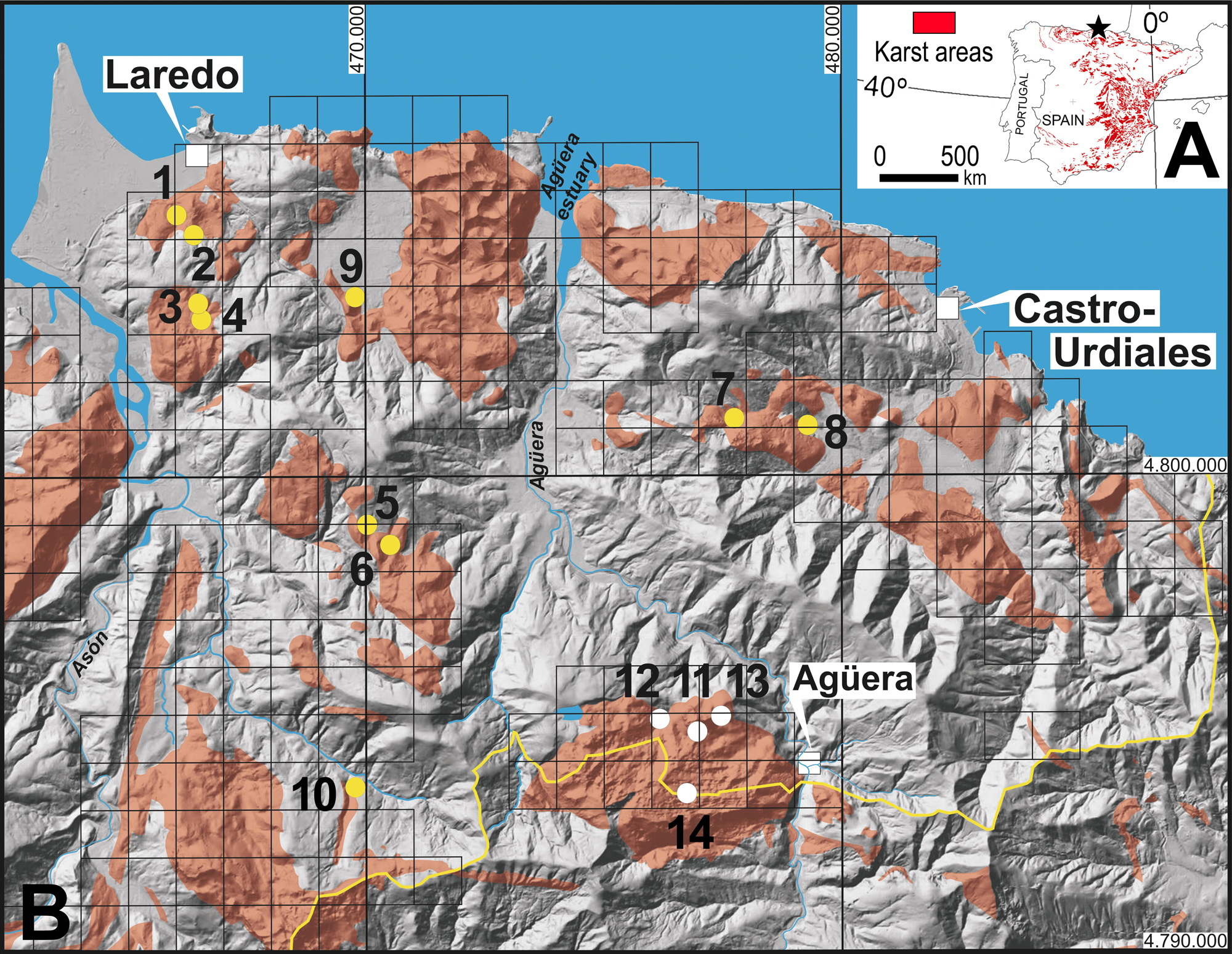
Type locality and habitus of *Ischyropsalis impressa* sp. nov. A: Entrance to La Baja Cave (arrow); B: Digital orthophoto with projection of plan view of the studied cave (surveyed August 2004-2006 by C.G. Luque and J. Ruiz-Cobo; see Ruiz-Cobo *et al*. 2008) and sampling sites of the new specie (star); C: Female in situ (La Baja Cave (CGL 269)), and D: Female in situ (Covacho del Cogorio Spring Cave (CGL 270)). Photo Collage by C.G. Luque. The final image courtesy of E. Peradalta.

**Figure 5.**
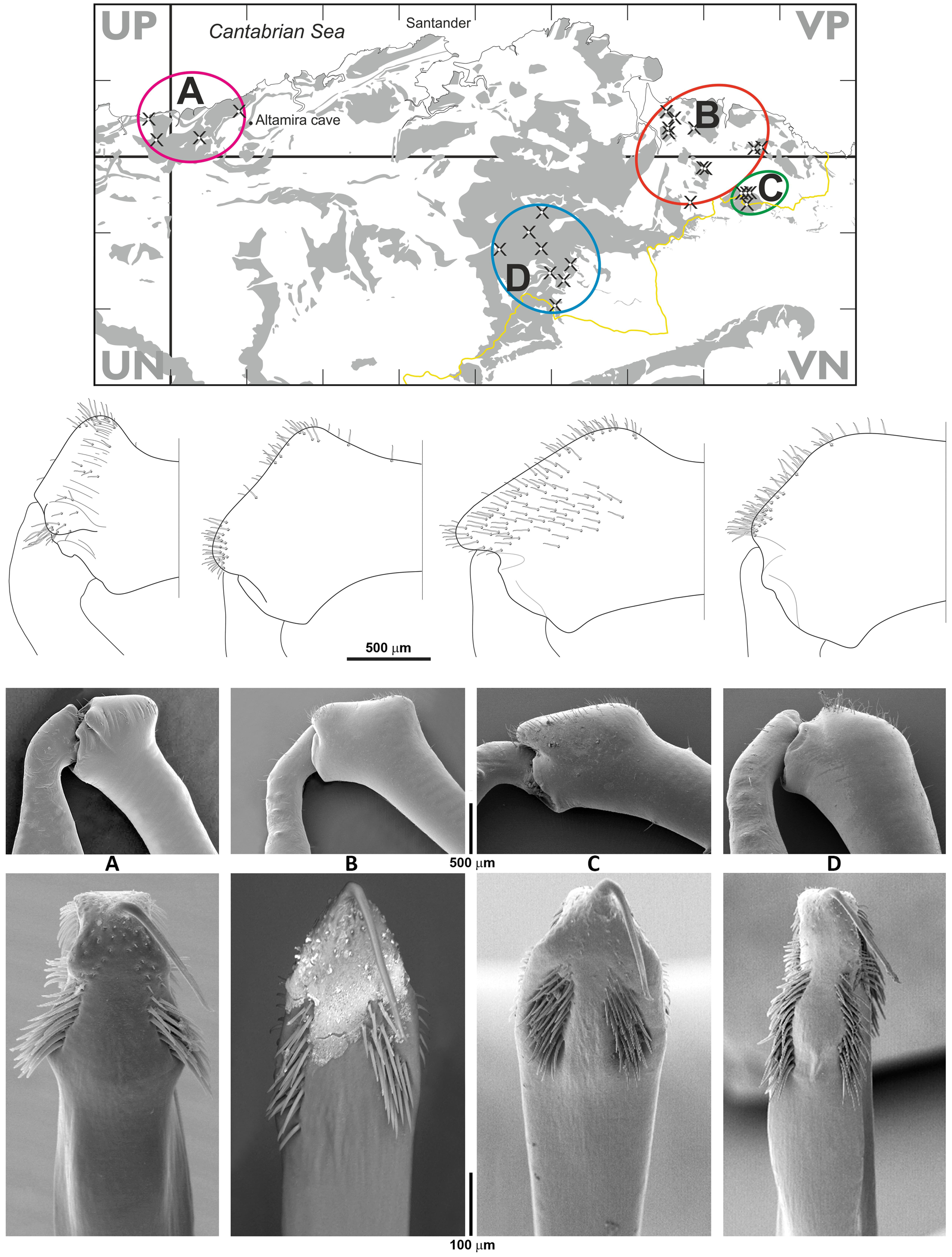
*Ischyropsalis impressa* sp. nov., male holotype from La Baja Cave: prosoma, lateral view (A); distal part of male chelicerae (medial view), apophyses and cheliceral secretion-extruding-field indicated by bristle-field (B). Chelicera of male paratypes from Hoyo de las Rebolligas Cave (C), Hoyo Molino Cave (D). Chelicera of female paratypes from La Baja Cave (E), Hoyo Menor-Rejuyo Cave (F), Hoyo de la Cubilla Cave (G). Total view of ventral penis (H). Arrows indicate diagnostic characters mentioned in the text.

**Figure 6.**
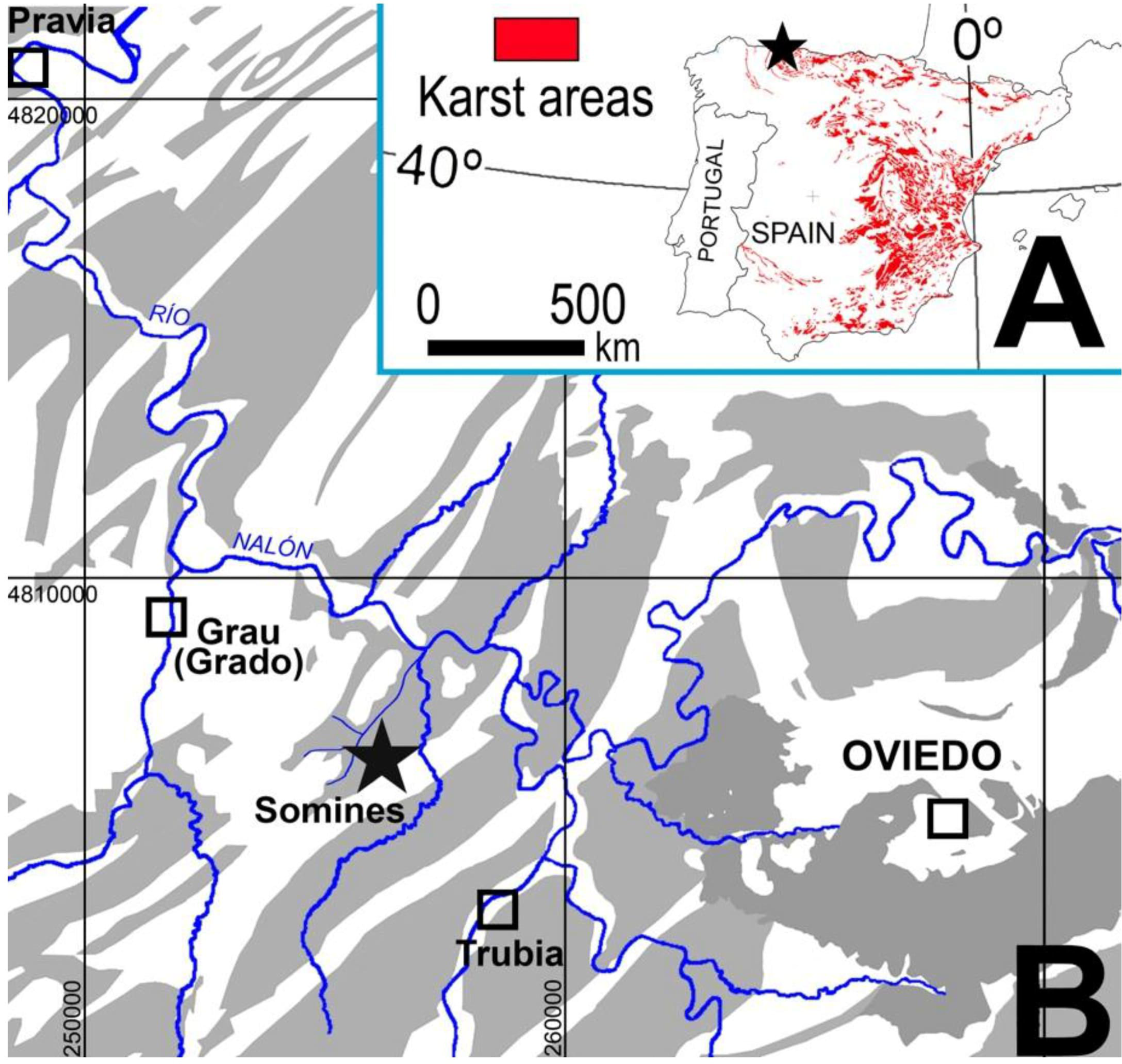
*Ischyropsalis impressa* sp. nov. from La Baja Cave (see figs. 38–41 in Luque and Labrada, 2012). A: Distal end of basal article with basal part of second article of male chelicera (MNCN 20.02/12858), medial view; B–C: Distal part of penis with glans and stylus (MNCN 20.02/17331): dorso-lateral view (B), dorsal view (C), and ventral view (C).

**Figure 7.**
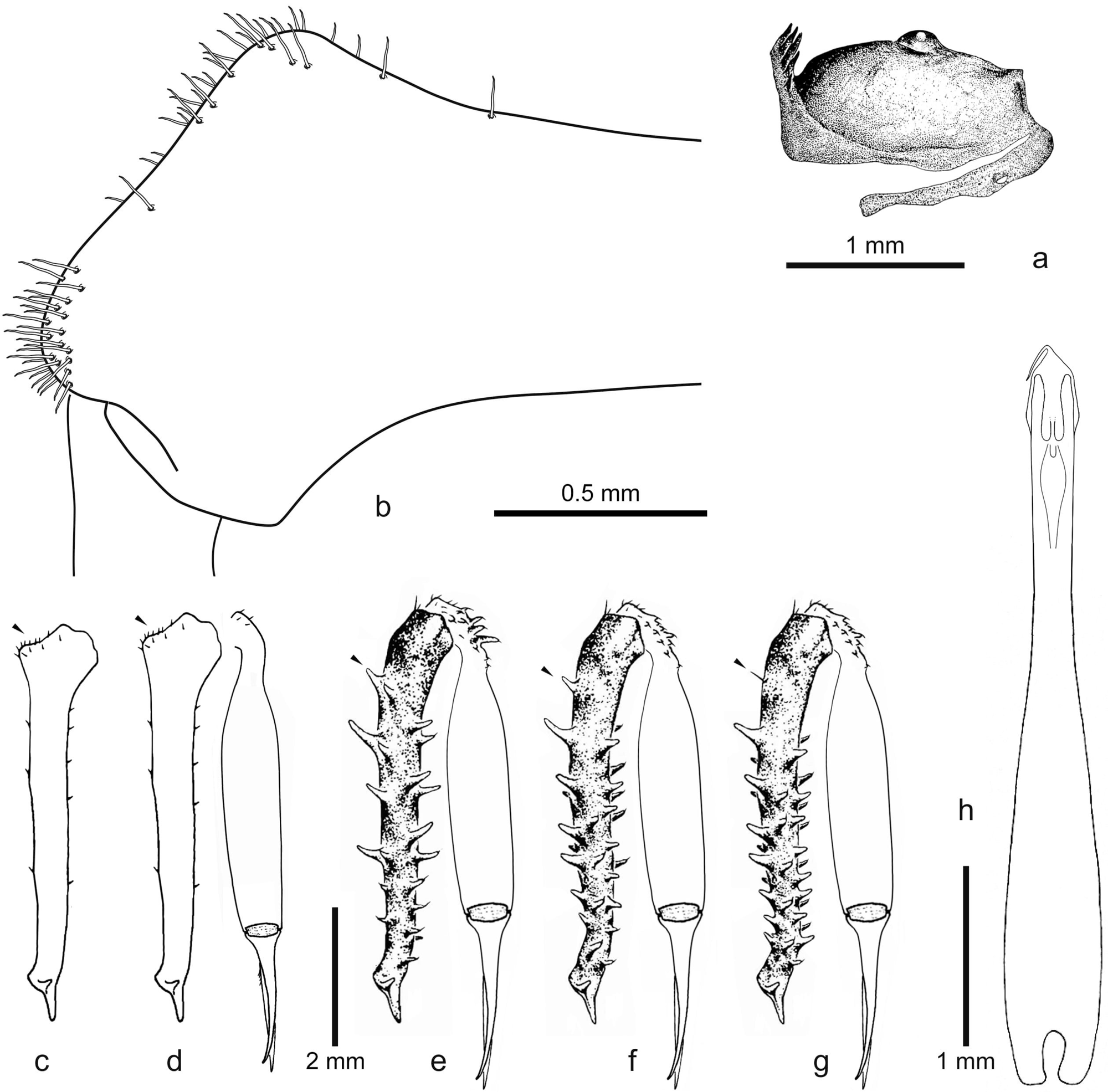
Type locality and habitus of *Ischyropsalis aguerana* sp. nov. A: Entrance of the caves of Hoyo de las Fuentes (*left*) and La Pasada (*right*); B: Digital orthophoto with projection of plan view of the studied cave (surveyed August 2004-2006 by C.G. Luque, E. Muñoz and A. Subiñas), the sampling of the new specie (star) and cave entrance (yellow dot); C: Alive female from La Pasada Cave (CGL 469). Collage of photographs by C.G. Luque.

### Etymology

The name of the species comes from the Latin word *impressio*, referring to Impress, a Santander-based consulting firm, where taxonomic studies were performed.

### Type material Holotype

SPAIN – Cantabria Province • ♂; Laredo; La Baja Cave; 30TVP6593705274, 50 m a.s.l.; 22 Aug, 2004; L. Labrada and C.G. Luque, leg.; MNCN 20.02/17329.

### Allotype

SPAIN – Cantabria Province • ♀; Laredo; La Baja Cave; 30TVP6593705274, 50 m a.s.l.; 22 Aug, 2004; L. Labrada and C.G. Luque, leg.; MNCN 20.02/17332.

### Paratypes (only adult specimens)

SPAIN – Cantabria Province • 3 ♂♂, 2 ♀♀; same collection data as for holotype and allotype; 22 Aug, 2004; in five vials with MNCN codes 20.02/12858, 20.02/17330, 20.02/17331, 20.02/17334, 20.02/17335.

### Additional material studied

SPAIN – Cantabria Province • 1 ♂, 1 ♀; Laredo; La Baja Cave; 12 Aug, 1985; CGL • 5 ♂♂, 4 ♀♀; Laredo; La Baja Cave; 28 Mar, 1992; CGL • 3 ♂♂, 3 ♀♀; Laredo; La Baja Cave; 22 Aug, 2004; CGL • 3 ♂♂, 5 ♀♀; Laredo; La Baja Cave; 28 Jul, 2006; CGL • 1 ♀ juv.; Liendo; Iseca Nueva; Covacho del Cogorio Spring Cave; 30TVP6978503552, 22 m a.s.l.; Aug, 1985; CGL • 1 ♀; Limpias; Seña; Perelada Cave; 30TVP6638403410, 180 m a.s.l.; 26 Aug, 1988; CGL • 1 ♂ juv.; Limpias; Seña; Hoyo de Peñaflor Cave; 30TVP6641104890, 95 m a.s.l.; 26 Dec, 1992; CGL • 1 ♂, 1 ♀; Limpias; Seña; Hoyo Molino Cave; 30TVP6650703093, 175 m a.s.l.; 31 Apr, 1988; CGL • 1 ♀; Limpias; Seña; Hoyo Molino Cave; 30TVP6650703093, 175 m a.s.l.; 31 Apr, 1993; CGL • 1 ♀; Laredo; La Baja Cave; 28 Mar, 1991; C.G. Luque, leg.; CJM 3756 • 1 ♂, 1 ♀; Laredo; La Baja Cave; 28 Mar, 1992; C.G. Luque, leg.; CRBA 97705 • 1 ♀; Guriezo; Hoyo Menor-Rejuyo Cave; 30TVN7053098550, 270 m a.s.l.; 6 Mar, 1988; C.G. Luque, leg.; CJM 3757 • 1 ♀; Castro-Urdiales; Sámano; La Lastrilla Cave; 30TVP7935500970, 60 m a.s.l.; 26 Jun, 1988; C.G. Luque, leg.; CJM 3758 • 6 ♀♀; Limpias; Hoyo Molino Cave; 3 Sep, 2010; L. Labrada & C.G. Luque, leg.; CGL • 1 ♂; Guriezo; Hoyo Menor-Rejuyo Cave; 9 Dec, 2010; L. Labrada & C.G. Luque, leg.; CGL • 1 ♀; Guriezo; Hoyo Menor-Tocinos Cave; 30TVP7018598775, 290 m a.s.l.; 31 Mar, 2013; L. Labrada & C.G. Luque, leg.; CGL • 1 ♀; Castro-Urdiales; Hoyo de la Cubilla Cave; 30TVP7770701185, 170 m a.s.l.; 31 Mar, 2013; L. Labrada & C.G. Luque, leg.; CGL • 1 ♂; Rasines; Hoyo de las Rebolligas Cave; 30TVN6959893325, 180 m a.s.l.; 9 Aug, 2013; C.G. Luque, leg.; CGL. Seven other adult females were captured by the same collectors as the holotype, and used for DNA extraction and sequencing with voucher numbers: BOS-OPI-00002 (2 ♀♀; Laredo; La Baja Cave; 24 Jul, 2012) (2 ♀♀; Limpias; Hoyo Molino Cave; 3 Sep, 2010), (2 ♀♀; Castro-Urdiales; La Lastrilla Cave; 31 Mar, 2013), (1 ♀; Guriezo; Hoyo Menor-Tocinos Cave; 31 Mar, 2013). Extracted specimens and DNA aliquots deposited in the Departments of Zoology at Oviedo.

### Remarks

The material originally collected between 1985 and 1992 by Luque (1991, 1992) from La Baja, Hoyo de Peñaflor, Hoyo Molino and Perelada Caves, incorrectly identified as *I. dispar*, as well as the material collected in the Hoyo Menor-Rejuyo and La Lastrilla Caves, incorrectly identified as *I. magdalenae*, are here reassigned to *I. impressa* sp. nov.

### Description

PROSOMA. Low prosoma, height at the eye area about 0.85 of its length. Fine and densely roughened, brown to black, slightly ascent from the border of the second thoracic tergite, and this tergite is ornamented with a transversal row of bristles with ten or more denticles of different sizes. Eye mound weakly developed. Eyes widely separated with small lenses (Figs. 4C-4D, 5A).

OPISTHOSOMA. Sclerotised brownish areas, finely granulated and surrounded by transversal grooves among shallow tergites. Each tergite with transverse rows of inconspicuous small tubercles, each one carrying a thick and short hair. Males and females with *scutum parvum* (Fig. 4C-4D).

CHELICERAE. Sexually dimorph. Short and slender, brown to black (can be up to ca. twice body length). Male basal article gracile and smooth (Figs. 5C-5D, 17G) with only a few short hairs, which widens distally to a weakly pronounced bulge (Figs. 5C–5D). The distal end of the basal article is slightly and ventrally angled, dorsally extended to a rounded apophysis (with a small glandular hairy area; Fig. 5B) and sloping flat to the insertion site of the distal article (medial view). Distal article is almost smooth, with only a few short hairs and rounded protuberances and bristles-like usually short and blunt set into a basal socket (see Luque & Labrada 2012, fig. 34).

PENIS. Total length of the penis 4.0 mm. Penis truncus penis (Fig. 5H) from base to below glans gradually narrowing. Slightly widening below glans, not dorsally bulged (dorso-lateral and dorsal views, Figs. 6B–6D) but a rather straight prolongation of the remainder of the truncus. Glans with conical apex gradually narrowing into a remarkably long stylus. Glans are separated into two sclerotised areas: dorsally with numerous setae arranged into two bands with 21 setae each (Fig. 6C). It ventrally displays short and narrow sclerite, almost parallel-sided. The basal end is divided into two lobes, each bearing long and abundant but loosely distributed bristles, which become absent towards the distal part of the glans or are reduced to only minute setae (Fig. 6D).

LEGS. Light brown, slender, medium length and all segments round in cross section. Metatarsus and tarsus in lighter colour. Base of femora whitish to yellowish, just as in *I. cantabrica* (see Luque & Labrada 2012, figs. 18–24). Smooth femora without sculpture elements (spines or tubercles), only with *microtrichia* and *setae*. *Trochanter* and *coxa* with several types of sculpture elements: granules, tubercles, cones and long *macrotrichia*, which arise from a basal socket.

MEASUREMENTS (mm) were taken for the holotype, allotype and paratypes (11 ♂♂, 6 ♀♀). Body length: holotype 5.04, ♂♂ paratypes 5.04–5.48, allotype 5.56, ♀♀ paratypes 5.56–5.97; length of basal *chelicerae* segment (paratypes in parentheses): holotype 4.76 (4.76–5.10), allotype 5.65 (5.65–5.72). Females show a wider range in body length, probably due to different stages of gravidity but their chelicerae are less variable in size than in male. Total length of leg II: 42.78 (42.78–44.32), 42.80 (42.80–44.21). Length of the leg II segments: femur: 8.88 (8.88–9.05), 9.41 (9.41–9.75); *patella*: 1.37 (1.37–1.42), 1.51 (1.51–1.57); tibia: 8.44 (8.44–8.98), 8.49 (8.49–8.86); *metatarsus*: 13.61 (13.61–13.98), 13.34 (13.34–13.73); *tarsus*: 10.48 (10.48–10.89), 10.05 (10.05–10.30). The measurements of the male (holotype) and female (in parentheses) *femora* from the first to the fourth pair of legs are 6.49 (6.91), 8.88 (9.41), 5.38 (5.61), 7.13 (7.42).

### Female description

CHELICERAE. Basal article more robust and armed with numerous long spines of different sizes over its whole length (7-8 longer spines on dorsal surface, 5-6 ventro-medially, and 6-8 ventro-laterally; Figs. 4C-4D, 5E–5G).

OVIPOSITOR. Bilobulate, with the ventral *apex* covered with few *setae*. The seminal receptacle has four short finger-like *ampullae*. In this respect, it does not differ from other *Ischyropsalis* species in the region (see Dresco 1968, fig. 19; Martens 1969b, figs. 66, 68).

### Variability

There seems to be no extraordinary variation, despite the general differences in *Ischyropsalis* species within the *dispar*-group (Luque & Labrada, 2012), e.g. female spination is more developed in larger specimens (see Martens 1969b: 222–224). The body is sometimes depigmented but *scutum parvum* uniformly dark brown. Only the population located in the eastern end (e.g. in La Lastrilla Limestone Cave system) is characterized by a body and legs with reduced pigmentation and small eyes.

### Distribution

Endemic to the northeastern coastal region of Cantabria. Restricted to a series of isolated karst areas and cave systems in the north foothills of the Cantabrian range. This area is formed by an Aptian Cretaceous limestone rock-type, called Urgonian limestone, with a radius of less than 15 km^2^ (Figs. 1-2, 17G).

### Ecology

The present records suggest that *I. impressa* sp. nov. is a troglobiont species. It has only been recorded in deep cave environment which is inhabited due to its cool, moist and dark microclimate. All karst areas and cave systems (Hoyo de Peñaflor, Hoyo Molino, Hoyo Menor, Hoyo de la Cubilla, La Lastrilla, La Baja, Covacho del Cogorio, etc.) are found with direct access to a subterranean river (the unsaturated or vadose zone of underground).

### *Ischyropsalis aguerana* sp. nov. Luque & Labrada, 2025

ZooBank taxon LSID: 14D2664B-3D2E-411A-9391-69167C34BD10 Figs. 1, 7-9, 17D

Diagnosis

A species of the westernmost *dispar*-group (see Luque & Labrada 2012) with long and thick *chelicerae*, which can be told apart by morphological characters and by allopatric range. Males are discernible by large distal apophyses on basal cheliceral article, with an extended bristle area (Figs. 8B–8C, 9A) and their distinct penis morphology (Figs. 8E, 9B–9D). Females are discernible by the presence of five large dorsal spines in the basal *chelicerae* article (Figs. 7C, 8D, see arrows).

**Figure 8.**
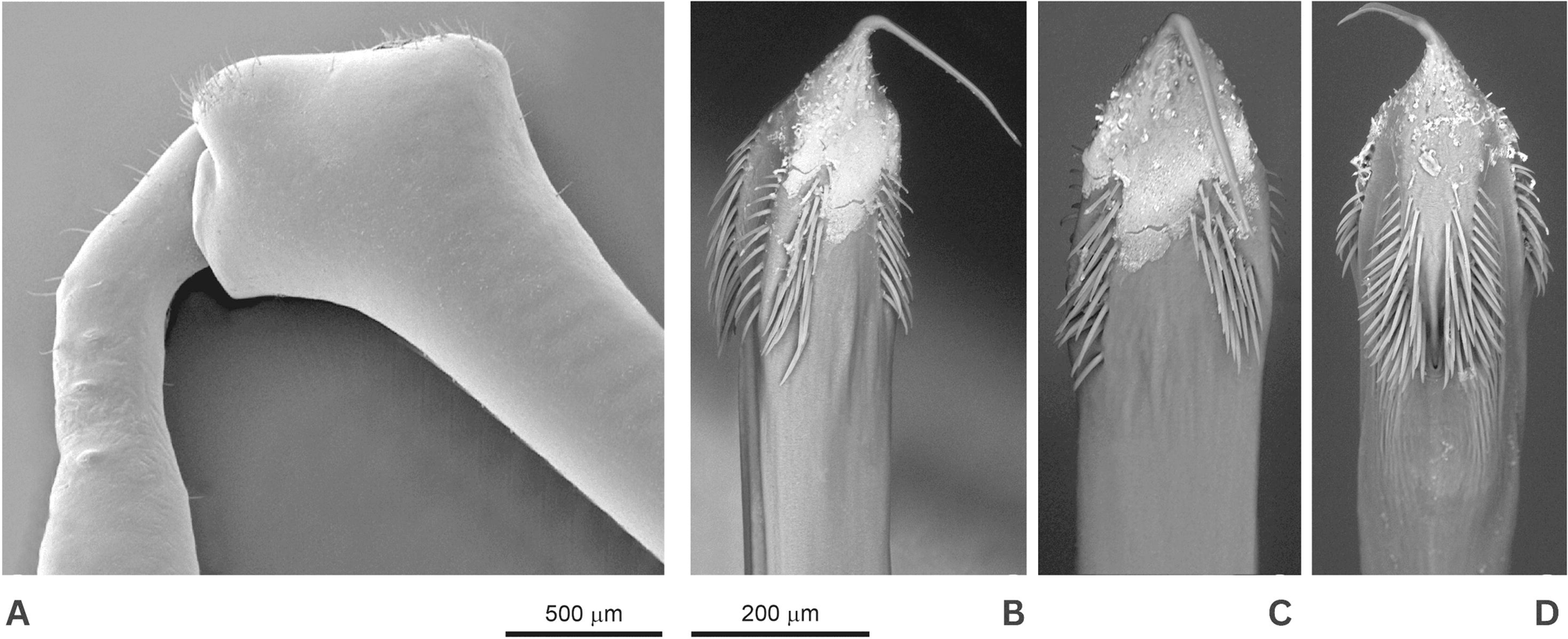
*Ischyropsalis aguerana* sp. nov., male holotype from La Pasada Cave: prosoma, lateral view (A); distal part of male chelicerae (medial view), apophyses and cheliceral secretion-extruding-field indicated by bristle-field (B). Chelicera of male (C) and female (D) paratypes from Hoyo de las Fuentes Cave. Penis ventral view (E). Arrows indicate diagnostic characters mentioned in the text.

**Figure 9.**
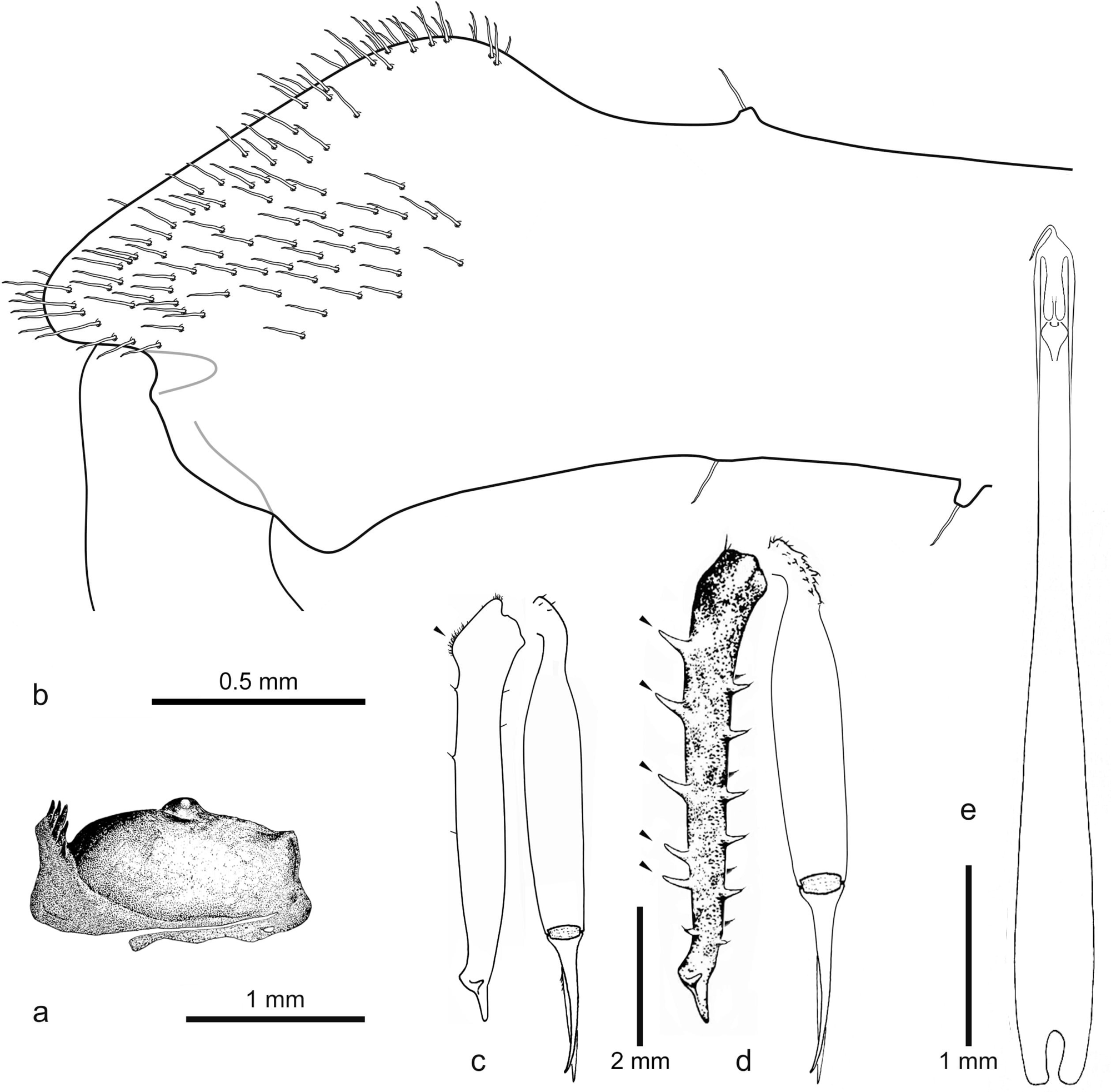
*Ischyropsalis aguerana* sp. nov. from La Pasada Cave, MNCN 20.02/12859 (type locality, Agüera, Guriezo, Cantabria; A-D; see also figs. 54–57 in Luque and Labrada 2012), and *Ischyropsalis dispar* Simon, 1872 from Albia Cave, MNCN 20.02/17337 (type locality, 30TVN9396857340, 810 m a.s.l., Villalba de Losa, Burgos, 19-VII-2008, leg. C.G. Luque & L. Labrada; E–H). Male cheliceral apophysis on distal end of basal article with basal part of second article of right chelicera (medial view) and cheliceral secretion-extruding-field indicated by bristle-field (A, E). Distal part of penis with glans and stylus: dorsal view (B, F), ventral view (C, G) and lateral view (D, H).

Etymology

The species name refers to the locality where it was found (Agüera).

Type material Holotype

SPAIN – Cantabria Province • ♂; Guriezo; La Pasada Cave; 30TVN7689794385, 440 m a.s.l.; 25 Apr, 1993; C.G. Luque, leg.; MNCN 20.02/17333.

Allotype

SPAIN – Cantabria Province • ♀; Guriezo; La Pasada Cave; 30TVN7689794385, 440 m a.s.l.; 22 Apr, 1993; C.G. Luque, leg.; MNCN 20.02/17336.

Paratypes (only adult specimens)

SPAIN – Cantabria Province • 1 ♂; same collection data as for holotype; MNCN 20.02/12859 • 4 ♂♂, 3 ♀♀; Guriezo; Hoyo de las Fuentes Cave; 30TVN7616294640, 320 m a.s.l.; Aug, 1985; C.G. Luque, leg.; misidentified as *Ischyropsalis dispar* by María Rambla (in seven vials with CRBA codes 97563, 97564, 97565, 97566, 97567, 97569,

97570).

Additional material studied

SPAIN – Cantabria Province • 1 ♂, 1 ♀; Guriezo; Hoyo de las Fuentes Cave; 30TVN7616294640, 320 m a.s.l.; 3 Aug, 2006; CGL • 1 ♀; Guriezo; Gueriza Cave (also is known as Agueriza or Agüeriza); 30TVN7669493073, 590 m a.s.l.; Oct, 1985; CGL • 1 ♀; Guriezo; La Tabla II Cave; 30TVN7739594717, 435 m a.s.l.; 25 Apr, 1993; CGL. Other adult female was captured by C.G. Luque and used for DNA extraction and sequencing, with voucher number BOS-OPI-00001 (1 ♀; Guriezo; La Pasada Cave; Aug, 2006). Extracted specimen and DNA aliquots deposited in the Department of Organisms and Systems Biology at Oviedo University.

Remarks

The material originally collected in August 1985 by C.G. Luque (unpublished material) from Hoyo de las Fuentes Cave, identified as *I. dispar* by M. Rambla (also the material collected in April 1993 from La Pasada Cave), is here reassigned to *I. aguerana* sp. nov.

Description

PROSOMA. Low prosoma, height at the eye area about 0.85 of its length. Fine and densely roughened, brown to black, slightly ascent from the border of the second thoracic tergite, and this tergite is ornamented with a transversal row of bristles with ten or more denticles of different sizes. Eye mound weakly developed. Eyes widely separated with small lenses.

OPISTHOSOMA. Brownish sclerotised areas, finely granulated, surrounded by transversal grooves among shallow tergites. Each tergite with transverse rows of inconspicuous small tubercles, each one carrying a thick and short hair. Males and females with *scutum parvum* (Fig. 7C).

CHELICERAE. Sexually dimorph. In general, long and slender, brown to black (can be up to ca. twice body length). Almost smooth male basal article, with a longitudinal row of 6-7 short piliferous tubercles, ventrally, and 2-3 short piliferous tubercles, dorsally (Fig. 30d). Male basal cheliceral article widens distally to a weakly pronounced bulge (Fig. 9A). The cheliceral apophysis is large, triangular and pointed with a rounded edge (lateral view), which harbours a large glandular and hairy area (Figs. 2C, 9A) bristle-like short and blunt, set into a basal socket (see Luque & Labrada 2012 fig. 34). The distal article with numerous and large piliferous tubercles rounded in shape at the dorsal knee, and some short hairs at the base of the movable finger.

PENIS. Total length of the penis 4.5 mm. Penis truncus (Fig. 8E) from base to below glans gradually narrowing. Below glans slightly widening, not dorsally bulged (dorso-lateral and lateral views, Figs. 9B-9D) but a rather straight prolongation of the remainder of the truncus. Glans with conical apex gradually narrowing into a remarkably long stylus. Glans are separated into two sclerotised areas: dorsally with numerous setae arranged into two bands with 35 setae each (Fig. 9B), and it ventrally displays long and narrow sclerite, almost parallel-sided. Basally end is divided into two lobes with long, abundant and dense bristles, thicker than in other parts of the glans, missing towards the distal part of glans or minute setae only (Fig. 9C). Differences between *I. dispar* and *I. aguerana* sp. nov. pennis morphology is shown in Fig 9E-H.

LEGS. Light brown, slender, medium length and all segments round in cross-section. Metatarsus and tarsus lighter in colour. Base of femora whitish to yellowish, and smooth femora without sculpture elements (spines or tubercles) only with microtrichia and setae. Trochanter and coxa with several types of sculpture elements: granules, tubercles and cones, and long macrotrichia, which arise from a basal socket.

Female description

CHELICERAE. Female basal article more robust and armed with numerous and long spines of different size over its whole length (5-6 longer spines on the dorsal surface, 6-7 ventro-medially, and 4-5 ventro-laterally; Figs. 7C, 8D).

OVIPOSITOR. Bilobulate, with the ventral apex covered with few setae. The seminal receptacle has four short finger-like ampullae. In this respect, it does not differ from other *Ischyropsalis* species in the region (see Dresco 1968 fig. 19; Martens 1969, figs. 66, 68).

MEASUREMENTS (mm) were taken for the holotype, allotype and paratypes (7 ♂♂, 7 ♀♀). Body length: holotype 5.14, ♂♂ paratypes 5.14–5.42, allotype 5.68, ♀♀ paratypes 5.68–5.75; length of basal chelicerael segment (paratypes in parentheses): holotype 5.30 (5.30–5.42), allotype 5.56 (5.56–5.85). Females show a wider range in body length, probably due to different stages of gravidity but their *chelicerae* are less variable in size than in male. Total length of leg II: 42.81 (42.81–43.67), 43.77 (43.77–44.87). Length of leg II segments: femur: 8.87 (8.87–9.03), 9.2 (9.2–9.37); *patella*: 1.37 (1.37–1.41), 1.52 (1.52–1.57); *tibia*: 8.39 (8.39–8.49), 8.59 (8.59–8.95); *metatarsus*: 13.65 (13.65–14.0), 13.81 (13.81–14.13); *tarsus*: 10.53 (10.53–10.74), 10.65 (10.65–10.85). The measurements of the male (holotype) and female (in parentheses) *femora* from the first to the fourth pair of legs are 5.81 (6.25), 8.87 (9.2), 5.2 (5.63), 6.9 (7.65).

Variability

There seems to be no extraordinary variation despite the general differences in *Ischyropsalis* species within the *dispar*-group (Luque & Labrada 2012), e.g. female spination is more developed in larger specimens (see Martens 1969b: 160–165).

Distribution

Endemic to the mountainous area of the middle Agüera basin. Only found in a particular karst area and cave system in the northern foothills of the Cantabrian range. Agüera area corresponds to thick Urgonian reef limestones from the Cretaceous, within a radius of less than 4 km^2^ (Figs. 1-2, 17D). Legal owner-ship of Agüera area, in dispute with neighbouring Biscay since 1552, is currently attributed to Cantabria since 2008 (Baró-Pazos 2010).

Ecology

The present records suggest that *I. aguerana* sp. nov. is a troglobiont species. It has only been recorded in deep cave environment which is inhabited due to its cool, moist and dark microclimate. Hoyo de las Fuentes is found with direct access to a subterranean river (the unsaturated or vadose zone of underground).

*Ischyropsalis damiani* sp. nov. López-Alonso, Pascual-Parra, Labrada, Luque, Cires, Arias & González-Toral, 2025

ZooBank taxon LSID: 589CFABA-618F-49CC-BCC1-B2C943302CAF Figs. 10-13

**Figure 10.**
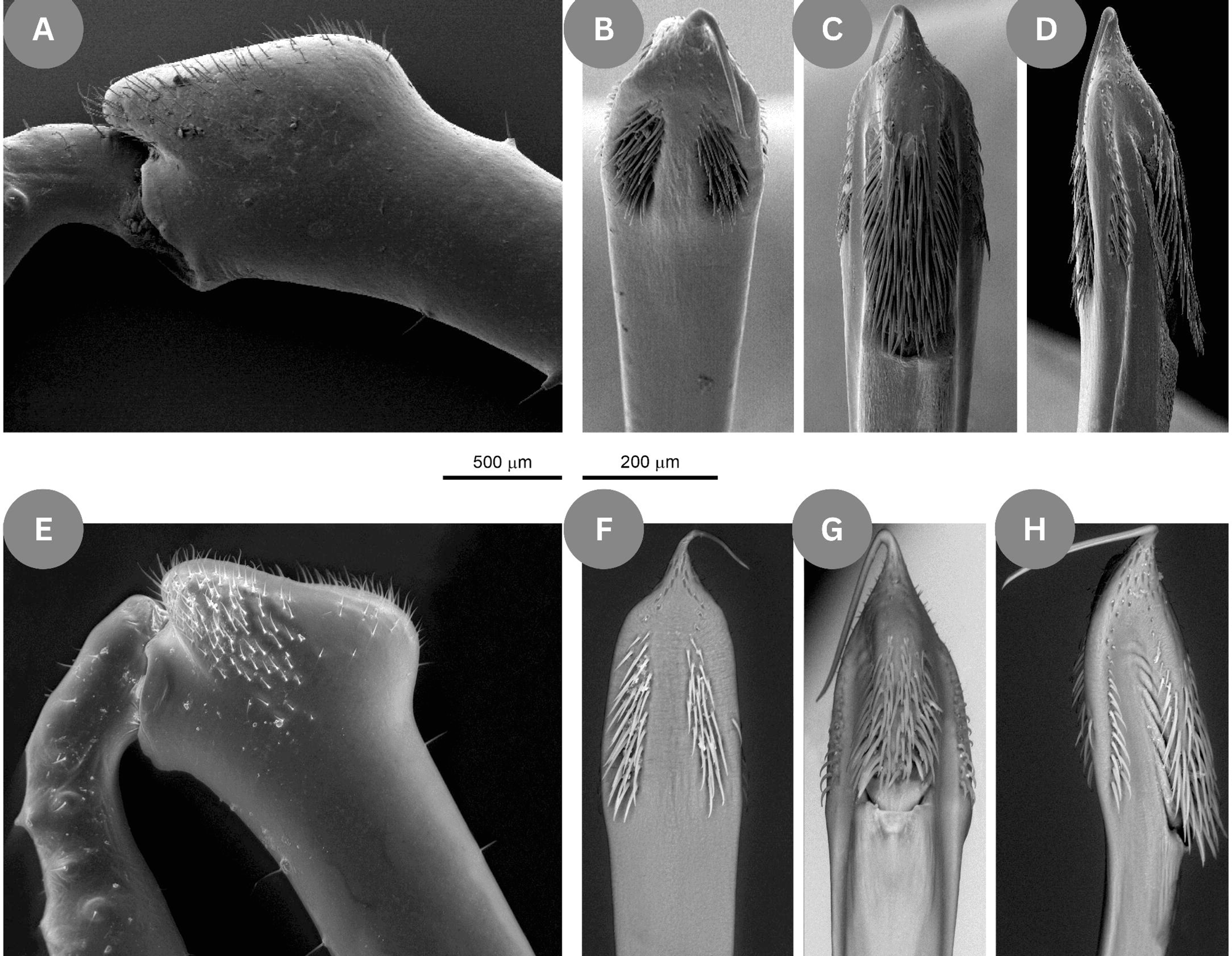
Type locality and habitus of *Ischyropsalis damiani* sp. nov. A: Entrance of the Elvira cave; B: Digital orthophoto with projection of plan view of the studied cave (surveyed August 2022 by López-Alonso, R and González-Toral, C.), the sampling of the new specie (star); C: Alive male from Elvira Cave.

Diagnosis

A species from the *nodifera*-group (see Martens 1969b) with short and slender *chelicerae*, which can be told apart by morphological characters and by allopatric range. Males are discernible by the distal flat pronounced bulge on basal cheliceral article (Fig. 11-12) mace-shaped distal end with a narrow bristle area (Fig. 11A) and by their distinct penis morphology (Fig. 11F-13). Females are discernible by the presence of short dorsal spines in the distal end of the basal article and the presence of several large dorsal and ventral spines in the basal cheliceral article (Fig. 11E).

**Figure 11.**
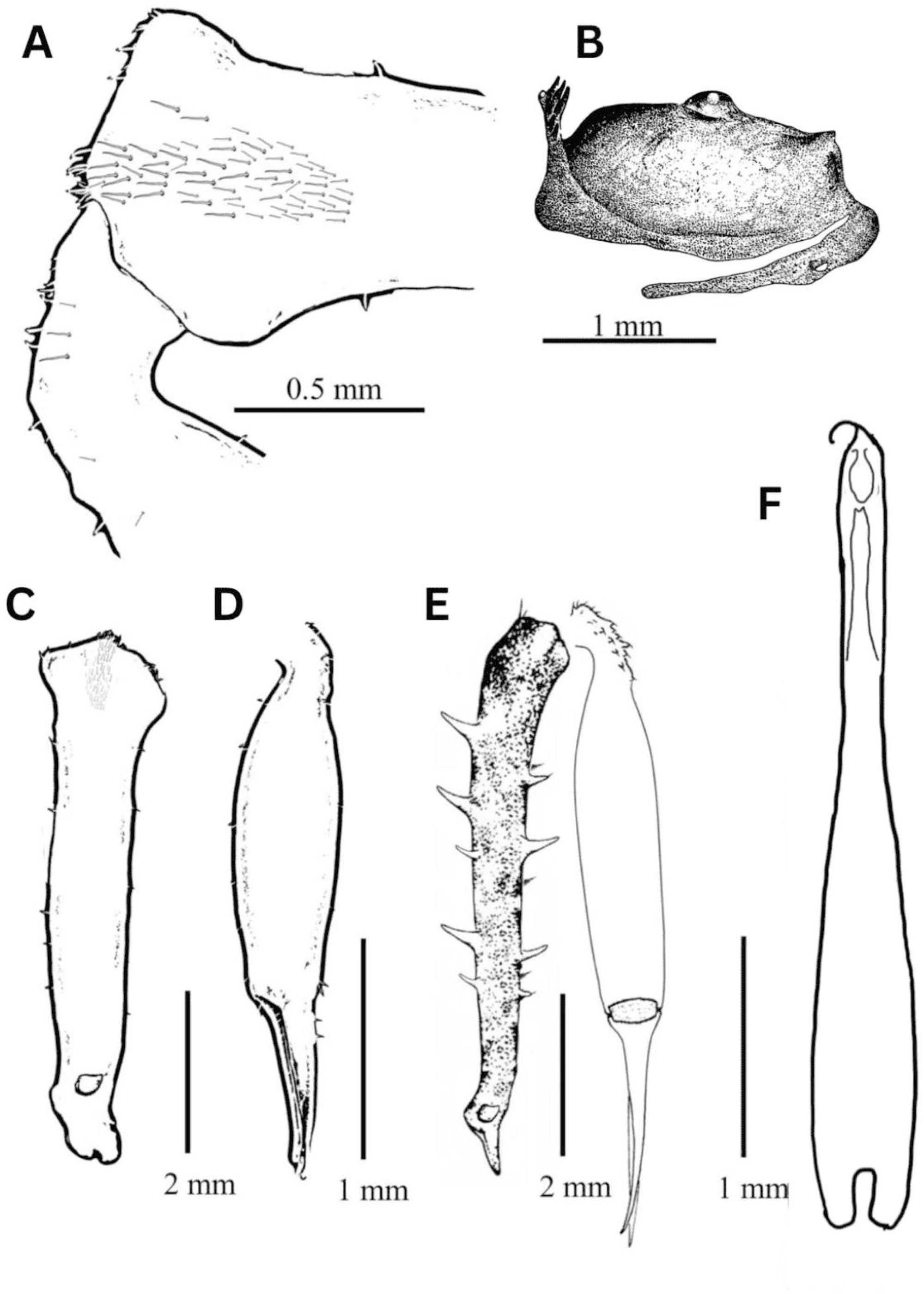
*Ischyropsalis damiani* sp. nov., male holotype from Elvira Cave: prosoma, lateral view (B); distal part of male chelicerae (medial view), apophyses and cheliceral secretion-extruding-field indicated by bristle-field (A). Chelicera of male (C-D). Chelicera of female (E). Total view of ventral penis (F).

**Figure 12.**
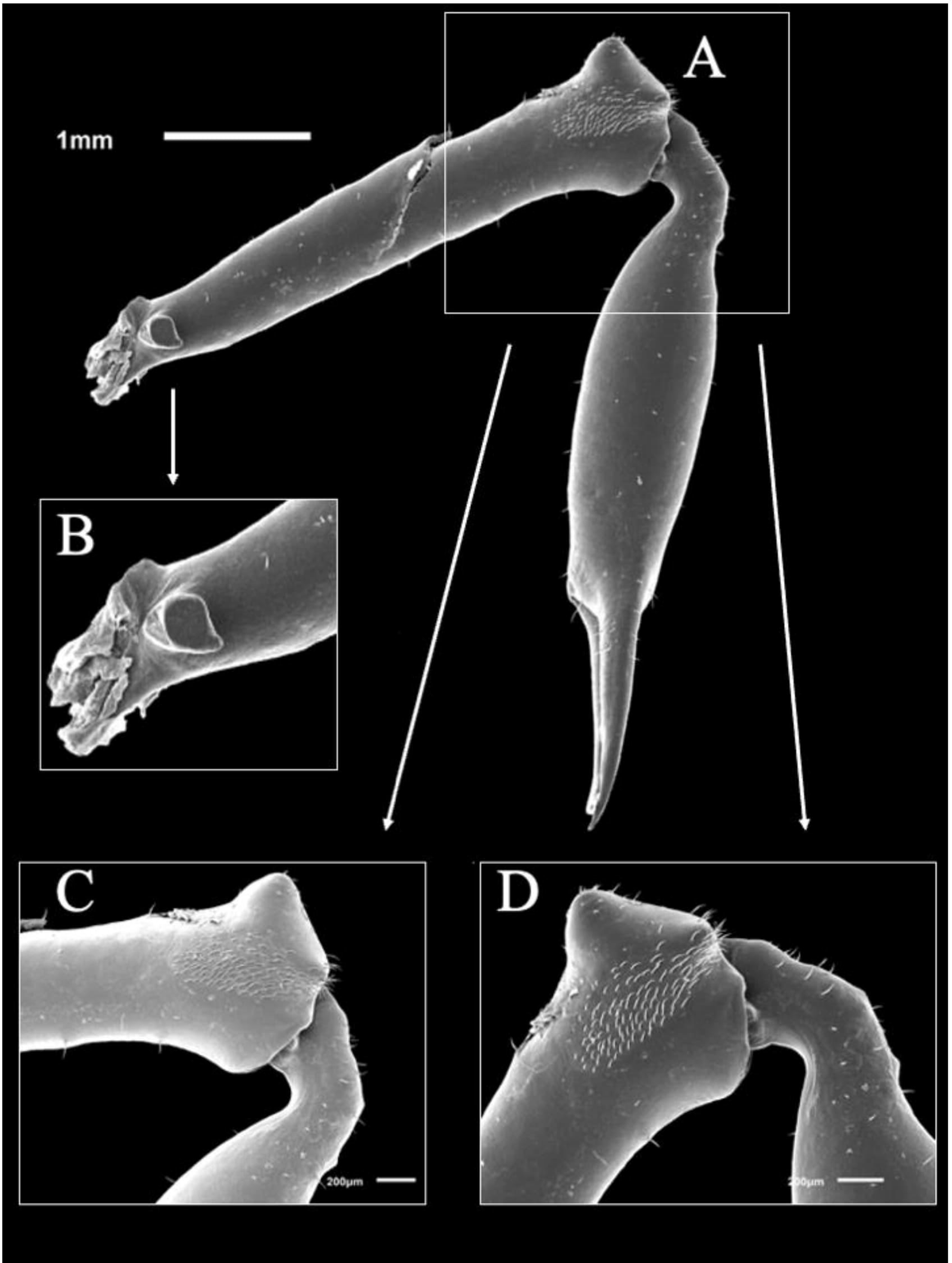
*Ischyropsalis damiani* sp. nov. from Elvira Cave (type locality, Somines, Grado, Asturias. A: Male chelicera. B: Proximal part of male chelicera. C-D: cheliceral secretion-extruding-field indicated by bristle-field.

**Figure 13.**
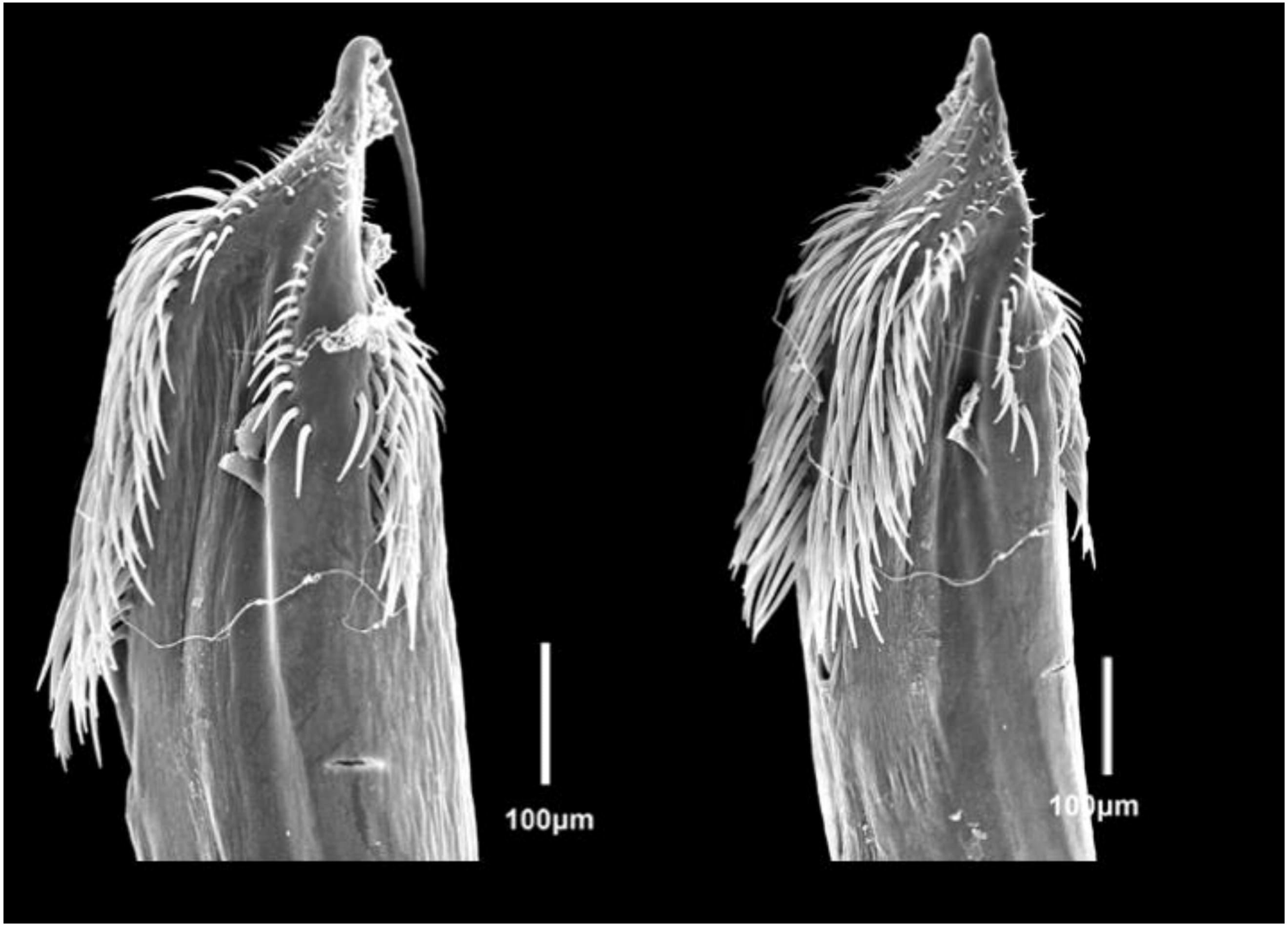
*Ischyropsalis damiani* sp. nov. from Elvira Cave (type locality, Somines, Grado, Asturias) Distal part of penis with glans and stylus: frontal and lateral view.

Etymology

The name of the species is a recognition to Damián’s friendship with the first author. The new species described herein is named after him.

Type material Holotype

SPAIN – Asturias Province • ♂; Somines, Grado; Cueva Elvira; 29T 41632 6161, 100 m a.s.l.; 02 Oct, 2023; López-Alonso R., González-Toral., Pascual-Parra E., leg.; BOS-OPI-00007.

Allotype

SPAIN – Asturias Province • ♀; Grado; Cueva Elvira; 29T 41632 6161, 100 m a.s.l.; 02 Oct, 2023; López-Alonso R., González-Toral., Pascual-Parra E., leg.; BOS-OPI-00008.

Paratypes (only adult specimens)

SPAIN – Asturias Province • 9 ♂♂, 3 ♀♀; same collection *data* as for holotype and allotype; 02 Oct, 2023; in 11 vials with the codes of the collection of Zoology of the University of Oviedo, from BOS-OPI-00003 to BOS-OPI-00007.

Description

PROSOMA. Low prosoma, height at the eye area about 0.8 of its length. Fine and densely roughened, grey to black, slightly ascent from border of second thoracic tergite. This tergite ornamented with a transversal row of bristles with ten or more denticles of different sizes. Eye mound weakly developed, eyes widely separated, with small lenses (Fig. 11B).

OPISTHOSOMA. Dark grey sclerotized areas, finely granulated, surrounded by transversal grooves between shallow tergites. Each tergite with transverse rows of inconspicuous small tubercles, each one carrying a thick and short hair. Males and females with *scutum parvum* (Fig. 10C).

CHELICERAE. Sexually dimorph. In general, short and slender, grey to black (can be up to ca. twice body length). Gracile and smooth male basal article (Fig. 11C-12B) with only few short hairs, which widens distally to a pronounced bulge (Fig. 11A, 12C-12D). Male basal cheliceral article widens distally to a weakly pronounced bulge (Fig. 12A-12D). The cheliceral apophysis is large, triangular and pointed with a rounded edge (lateral view), which harbours a medium-size and wide glandular and hairy area (Figs. 11A, 12A-12D) bristles-like short and blunt, set into a basal socket (see Luque and Labrada, 2012). The distal article with numerous and large piliferous tubercles, rounded shape at the dorsal knee (Figs. 11A, 12C-12D) and some short hairs at the base of the movable finger.

PENIS. Total length of the penis 4.8 mm. Penis truncus (Fig. 11F) from base to below glans gradually narrowing. Below glans slightly widening, not bulged dorsally (dorso-lateral and dorsal views, Fig. 13) but a rather straight prolongation of the remainder of the truncus. Glans with conical apex gradually narrowing into a long stylus. Glans separated into two sclerotised areas: dorsally with numerous setae (Fig. 13). The basal end divided into two lobes with long, abundant and dense bristles, with numerous setae towards the distal part of the glans (Fig. 13).

LEGS. Slender, medium length, all segments round in cross section. Light grey metatarsus. Yellowish tarsus and base of femora. Smooth femora without sculpture elements (spines or tubercles), only with microtrichia and setae. Trochanter and coxa with several types of sculpture elements: granules, tubercles and cones, and long macrotrichia, which arise from a basal socket.

Female description

CHELICERAE. Female basal article more robust and armed with numerous and long spines of different size over its whole length (3-5 longer spines on dorsal surface, 4-6 ventro-medially, and 3-4 ventro-laterally).

OVIPOSITOR. Bilobulate, with the ventral apex covered with few setae. The seminal receptacle has four short finger-like ampullae. In this respect it does not differ from other *Ischyropsalis* species in the region (see Dresco, 1968, fig. 19; Martens, 1969b, figs. 66, 68).

MEASUREMENTS (mm) were taken for the holotype, allotype and paratypes (9 ♂♂, 3 ♀♀). Body length: holotype 5.28, ♂♂ paratypes 5.33–6.03, allotype 6.54, ♀♀ paratypes 6.55–

6.99; length of basal *chelicerae* segment (paratypes in parentheses): holotype 3.84 (3.82–4.07), allotype 4.36 (3.71–4.46). Females show a wider range in body length probably due to different stages of gravidity, but their *chelicerae* are less variable in size than in male. Total length of leg II: 29.16 (28.24–33.08), 28.79 (27.73–30.58). Length of the leg II segments: femur: 6.81 (5.81–7.04), 6.04 (5.67–6.74); *patella*: 1.55 (1.38–1.71), 1.66 (1.55–1.64); *tibia*: 5.59 (5.15–5.95), 5.66 (4.96–5.95); *metatarsus*: 8.04 (7.57–8.92), 7.65 (7.53–8.49); *tarsus*: 7.55 (7.52–8.92), 7.44 (7.44–7.76). The measurements of the male (holotype) and female (in parentheses) *femora* from the first to the fourth pair of legs are 4.75 (4.32), 6.81 (6.32), 3.85 (3.52), 5.61 (5.11).

Variability

There seems to be no extraordinary intraspecific variation despite the general differences in *Ischyropsalis* species within the *nodifera*-group (Luque & Labrada 2012), e.g. female spination is more developed in larger specimens (see Martens 1969b: 222–224). The body is sometimes depigmented but *scutum parvum* is uniformly light grey. The differences with *Ischyropsalis asturica* Roewer, 1950, which is known from a single female specimen collected in Oviedo, Asturias, have also been taken into account. Although *I. asturica* is only represented by a single specimen, our material (n = 14) displays a denser cheliceral armature (with more spines) and a more slenderer cheliceral shape. Martens (1969 fig. 49h, vial with code SMF/RII/5042) synonymized *I. asturica* with *I. nodifera*, attributing several nominal species to the high variability of cheliceral morphology within *I. nodifera*. Given the well-known morphological similarity among *Ischyropsalis* females and the unresolved species boundaries within the genus, we have carefully evaluated these differences. In addition to the consistent morphological distinctions, our species is also genetically distinct from *I. nodifera*, further supporting its recognition as a new and separate species.

Distribution

Endemic to the central region of Asturias, restricted to an isolated karst area and cave in the central part of Asturias. This area is formed by a Carboniferous limestone rock-type, called Grey or Mountain limestone. (Fig. 3).

Ecology

The present records suggest that *I. damiani* sp. nov. is a troglobiont species. It has only been recorded in the cave environment which is cool, moist and dark. The cave has direct access to a subterranean river (the unsaturated or vadose area of the ground).

Phylogenetical analyses

The COI dataset consisted of 26 different taxa and 56 different *Ischyropsalis* sequences, with a length of 358 pb, 108 parsimonious-informative sites (Table 1), and the EF1-α dataset consisted of 21 different taxa and 41 different sequences, with a length of 607 pb, 133 parsimonious-informative sites (Table 1). The COIxEF1-α dataset consisted of 21 different taxa and 35 different sequences, with a length of 963 pb, 206 parsimonious-informative sites (Table 1).

The *COI* dataset analyses retrieved topologies (Fig. 14) with a highly supported *Ischyropasalis* clade (BS-ML:100; BI-PP:100) in which the following clades can be distinguished: the *I. hellwigii* exclusive clade group (BI-PP:51), the *I. carli* exclusive clade (BS-ML:100; BI-PP:100), the *I. dentipalpis*-group (BS-ML:99; BI-PP:100), and the Alpine-Iberian-*manicata* group (BS-ML:63; BI-PP:58). The latter was formed by four sister subclades: the *I. muellneri*-*I. strandii* subclade (BS-ML:96; BI-PP:84), the *I. ravasinii-I. lithoclasica-I. kollari-I. manicata-I. adamii-I. alpinula* subclade (BS-ML:54; BI-PP:64) formed by *I. ravasinii*-*I. lithoclasica*-*I. kollari* (BS-ML:98; BI-PP:89) and the *I. adamii*-*I. manicata-I. alpinula* subclade (BS-ML:63; BI-PP:64), and two Iberian-Cantabrian subclade, one comprising *I. pyrenaea-I. luteipes-I.* sp-*I. carli* subclade (BS-ML:92; BI-PP:83). *Ischyropsalis* carli JX563638, which is from the *manicata-*group, clustered within the Iberian-Cantabrian subclade. Another Iberian-Cantabrian subclade, formed by individuals from *I. hispanica*, *I. nodifera*, *I. cantabrica*, *I. damiani* sp. nov., *I. impressa* sp. nov. and *I. aguerana* sp. nov., *I. petiginosa*, *I. navarrensis*, *I. robusta*, *I. magdaleneae* and *I. dispar* (BS-ML:98; BI-PP:100) is also present. The Eastern Iberian subclade (as illustrated in Schönhofer et al. 2015) did not generate species-exclusive inner subclades. Similarly, Western Iberian subclade (as illustrated in Schönhofer et al. 2015) did not generate species-exclusive subclades, except for the highly-supported *I. dispar* inner subclade (BS-ML:98; BI-PP:82).

**Figure 14.**
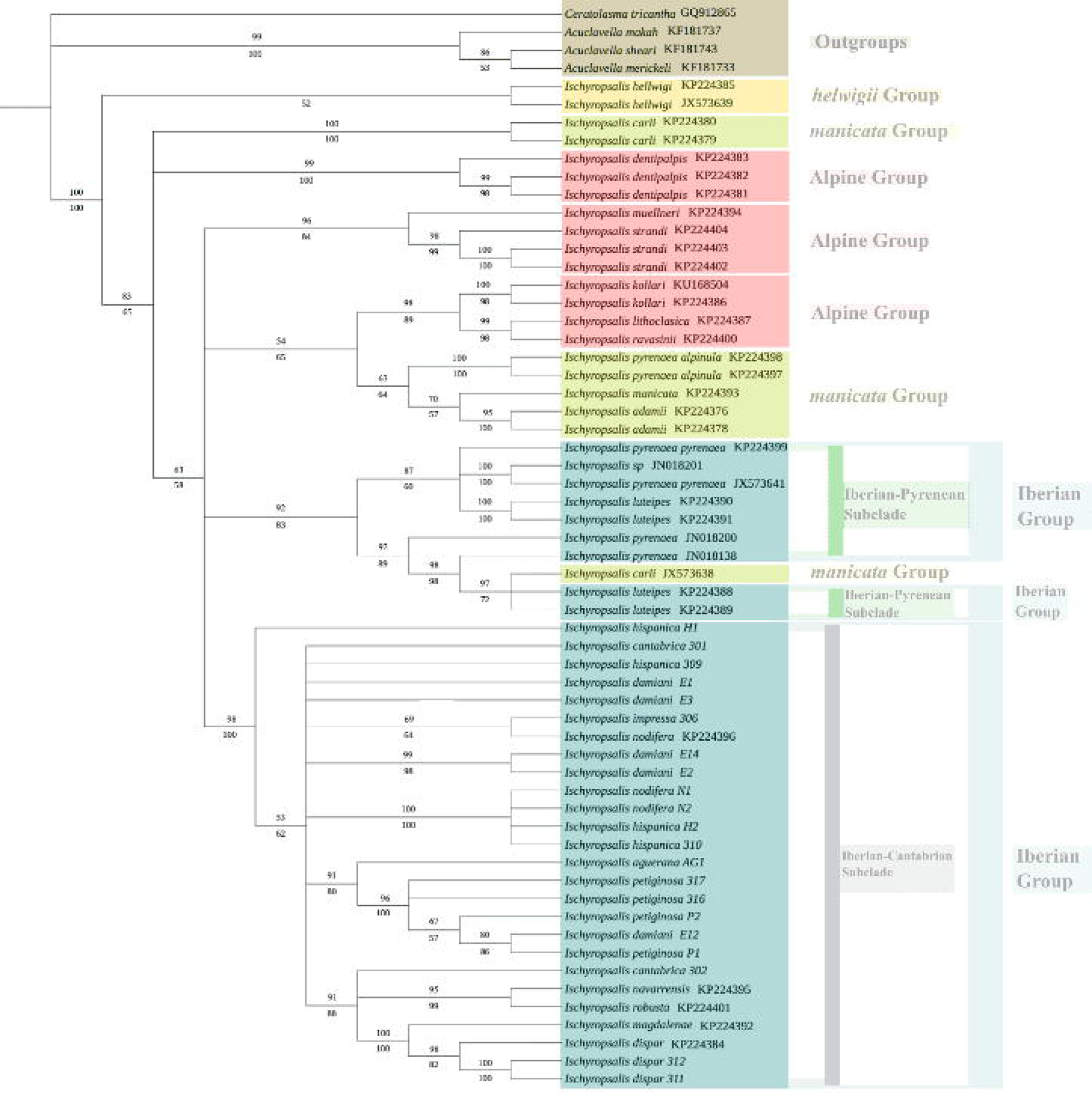
Consensus topology obtained from the Maximum Likelihood (ML) and Bayesian Inference (BI) analyses of the *COI* dataset. Values beneath the branches indicate the Posterior Probability (PP) support from the BI analysis, while values above the branches represent the ultrafast bootstrap (BT) support from the ML analysis.

The EF1-α analyses (Fig. 15) retrieved topologies also portrayed *Ischyropasalis* as a monophyletic clade (BS-ML:100; BI-PP:100). Within the *Ischyropasalis* clade, the following sublclades were found: the Iberian-Alpine clade (BS-ML:51; BI-PP:89), formed by *I. dentipalpis*, *I. strandi*, *I. ravasinii*, *I. lithoclasica*, *I. kollari*, *I. luteipes*, *I. pyrenaea*, *I. navarrensis*, *I. robusta*, *I. magdalenae*, *I. dispar*, *I. nodifera*, *I. petiginosa*, *I. hispanica* and *I. damiani* sp. nov; the *manicata*-group (BS-ML:92; BI-PP:98), comprising *I. adamii*, *I. carli* and *I. alpinula*, and the *hellwigii*-group, formed by *I. hellwigii* subsp. *hellwigii* and *I. hellwigii* subsp. *lucantei* (BS-ML:98; BI-PP:100). Within the Iberian-Alpine, the Iberian subclade (BI-PP:69) and the Alpine subclade (BS-ML:98; BI-PP:100) could also be distinguished. The weakly supported Iberian clade was formed by 2 sister subclades: the moderately supported Iberian Pyrenean subclade (BS-ML:54; BI-PP:90), formed by *I. luteipes* and *I. pyrenaea* and a highly supported Iberian Cantabrian subclade (BS-ML:96; BI-PP:100), comprising *I. navarrensis*, *I. robusta*, *I. magdalenae*, *I. dispar*, *I. nodifera*, *I. petiginosa*, *I. hispanica* and *I. damiani* sp. nov. Within the latter subclade, individuals from *I. damiani* sp. nov. (BS-ML:69; BI-PP:98), *I. hispanica* (BS-ML:92; BI-PP:56) and *I. dispar* (BS-ML:95; BI-PP:65) formed species-specific clades. The *I. petiginosa* sample has a moderate to highly-supported sister relationship with the *I. hispanica* exclusive clade (BS-ML:74; BI-PP:97). Similarly, the *I. magdalenae* individual was closely related to the *I. dispar* exclusive clade with high support (BS-ML:100; BI-PP:100). On the other hand, the *I. nodifera* sequenced individuals did not form any species-exclusive clade, generating two separate individual clades within the Western Iberian subclade.

**Figure 15.**
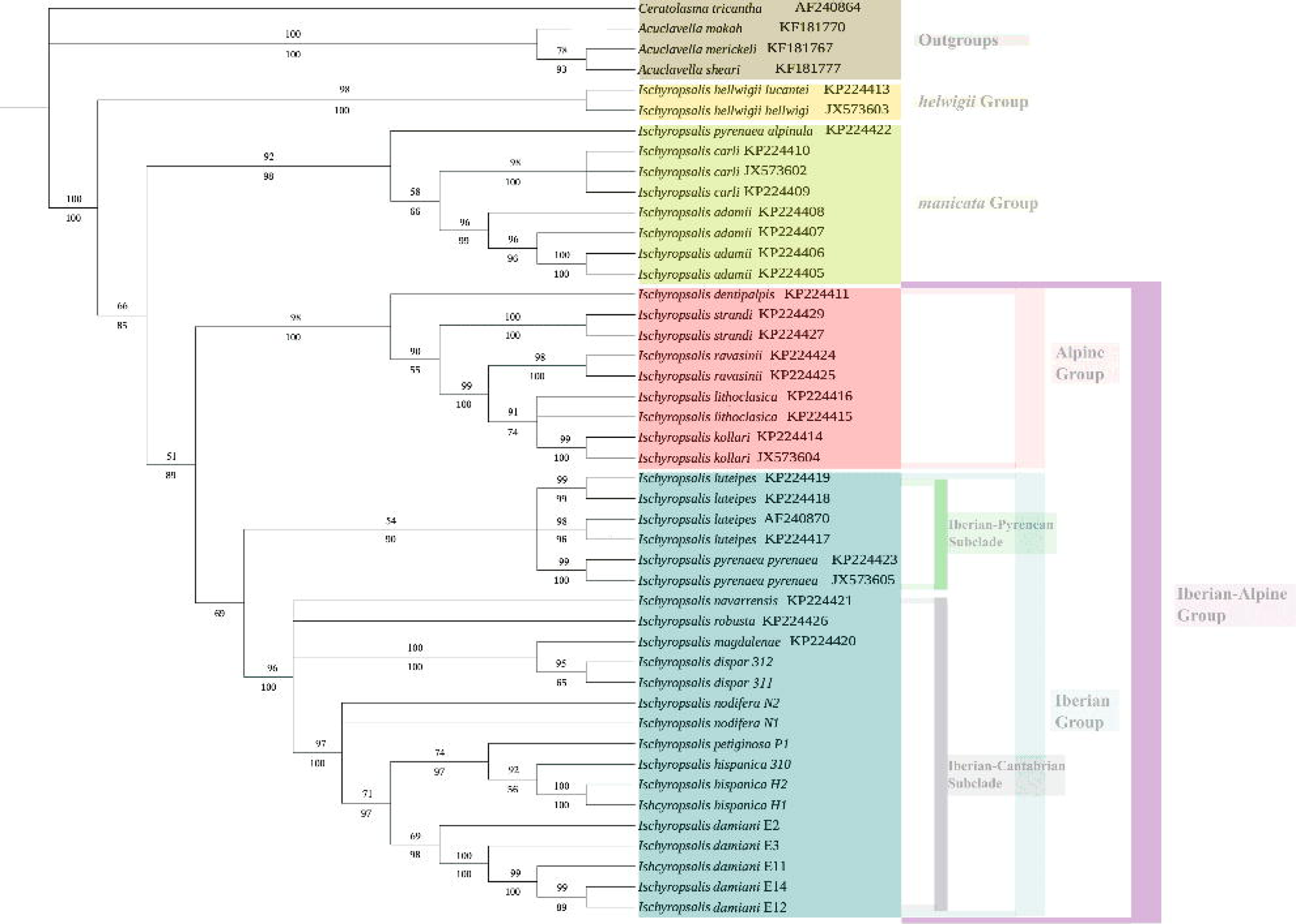
Consensus topology obtained from the Maximum Likelihood (ML) and Bayesian Inference (BI) analyses of the *EF1-α* dataset. Values beneath the branches indicate the Posterior Probability (PP) support from the BI analysis, while values above the branches represent the ultrafast bootstrap (BT) support from the ML analysis.

The EF1-α-COI analyses retrieved topologies (Fig. 16) in which *Ischyropasalis* was monophyletic (BS-ML:100; BI-PP:100). Within this topology the Iberian-Alpine clade (BS-ML: 74; BI-PP: 61), formed by, *I. dentipalpis*, *I. strandi*, *I. ravasinii*, *I. lithoclasica*, *I. kollari*, *I. luteipes*, *I. pyrenaea*, *I. navarrensis*, *I. robusta*, *I. magdalenae*, *I. dispar*, *I. nodifera*, *I. hispanica*, *I. petiginosa* and *I. damiani* sp. nov; the *manicata*-group (BS-ML: 87; BI-PP: 97), comprising *I. adamii*, *I. carli* and *I. alpinula*, and the basal *hellwigii*-group, formed by *I. hellwigii* subsp. *hellwigii* and *I. hellwigii* subsp. *lucantei* (BS-ML: 98; BI-PP: 100) could be found. Within the Iberian-Alpine, the Iberian subclade (BS-ML: 90; BI-PP: 78) formed by *I. luteipes*, *I. pyrenaea*, *I. navarrensis*, *I. robusta*, *I. magdalenae*, *I. dispar*, *I. nodifera*, *I. hispanica*, *I. petiginosa* and *I. damiani* sp. nov; and the Alpine subclade (BS-ML: 96; BI-PP: 98), comprising *I. dentipalpis*, *I. strandi*, *I. ravasinii*, *I. lithoclasica* and *I. kollari*, could also be distinguished. Within the moderately supported Iberian subclade, the strongly supported Iberian Pyrenean subclade (BS-ML: 98; BI-PP: 97), formed by, *I. luteipes* and *I. pyrenaea* and the highly supported Iberian Cantabrian subclade (BS-ML: 100; BI-PP: 100), comprising *I. navarrensis*, *I. robusta*, *I. magdalenae*, *I. dispar*, *I. nodifera*, *I. petiginosa*, *I. hispanica* and *I. damiani* sp. nov. Within the latter subclade, individuals from *I. damiani* sp. nov. (BI-PP: 100) and *I. dispar* (BS-ML: 98; BI-PP: 94) formed species-specific clades. The *I. petiginosa* individual has a moderate to sister relationship with the *I. damiani* sp. nov. exclusive clade (BS-ML: 88; BI-PP: 72). Similarly, the *I. magdalenae* individual was closely related to the *I. dispar* exclusive clade with high support (BS-ML:100; BI-PP:100). On the other hand, the *I. nodifera* sequenced individuals formed a species-exclusive clade with the *I. hispanica* (BI-PP:91). The other sequences of *I. hispanica* formed a species exclusive clade (BS-ML: 81; BI-PP: 100).

**Figure 16.**
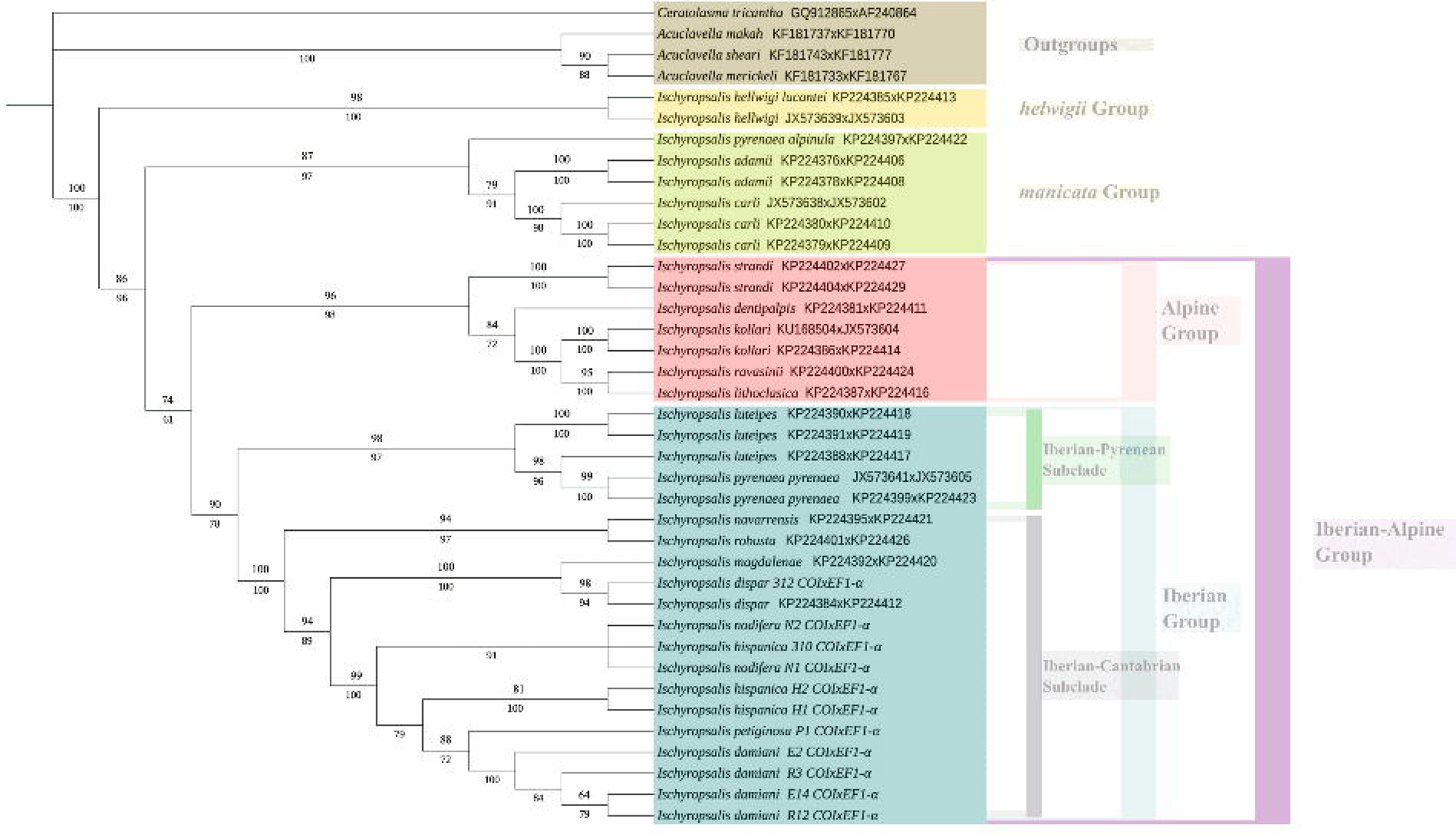
Consensus topology obtained from the Maximum Likelihood (ML) and Bayesian Inference (BI) analyses of the COIxEF1-α dataset. Values beneath the branches indicate the Posterior Probability (PP) support from the BI analysis, while values above the branches represent the ultrafast bootstrap (BT) support from the ML analysis.

## Discussion

The use of phylogenetic methods based on nuclear and mitochondrial markers (Schönhofer *et al*. 2015; Prieto *et al*. 2013), together with male genital and cheliceral morphology (e.g. Martens 1978; Schmidt *et al*. 2024) may clarify the taxonomy of *Ischyropsalis*. In this study, morphological data and molecular methods based on COI and EF1-α have provided a clearer taxonomy of the Cantabrian *Ischyropsalis* taxa. Our phylogenetic results revealed the existence of at least 25 valid species, 12 of which occur exclusively in the Iberian Peninsula, except *I. luteipes*, which also is found in the French Massif Central (Prieto *et al*. 2013) and including 3 new species (*I. damiani* sp. nov., *I. impressa* sp. nov. and *I. aguerana* sp. nov.). Our phylogenetic analysis (based on two of the three molecular markers used by Schönhofer *et al*. (2015)) reconfirmed the same clades, with the newly sequenced species placed within the Iberian Clade. Furthermore, our EF1-α and COIxEF1-α topologies also supported the existence of the Iberian Pyrenean subclade, which includes species occurring in the Pyrenean (with *I. pyrenaea* and *I. luteipes* form the Pyrenees and the French Massif Central), and the Iberian Cantabrian subclade, with species belonging to the *I. dispar*-group and *nodifera-*group our newly sequences species (*I. hispanica*, *I. cantabrica*, *I. petiginosa* and *I. damiani* sp. nov.). This placement supports the hypothesis of Schönhofer *et al*. (2015) who suspected that, based on morphological characters and distribution, *I. cantabrica* and *I. gigantea* likely belongs to the *dispar*-group.

We were able to generate COI sequences from three of these three new species, *I. damiani* sp. nov., *I. impressa* sp. nov. and *I. aguerana* sp. nov., which revealed the close phylogenetic relationship with other species occurring in the western Iberian Peninsula (*I. hispanica*, *I. nodifera*, *I. cantabrica*, *I. petiginosa*, *I. navarrensis*, *I. robusta*, *I. magdalenae* and *I. dispar*). In this sense, the morphological analysis of the Iberian taxa revealed that their cheliceral apophysis is generally large and often triangular and pointed with a rounded edge (lateral view), while the female *chelicerae* provided with numerous spines of variable size. Furthermore, a well-marked character in *I. impressa* sp. nov. and *I. aguerana* sp. nov. is a divided two-lobed sclerite of the glans. According to this morphological character, the new species from Cantabria (*I. aguerana* sp. nov. and *I. impressa* sp. nov.) would belong to the *dispar*-group sensu Schönhofer *et al*. (2015), which comprises the Iberian-Cantabrian taxa (*I. dispar*, *I. magdalenae, I. cantabrica, I. navarrensis* and *I. gigantea*), which is also supported by the phylogenetic position of *I. aguerana* sp. nov. and *I. impressa* sp. nov. in the COI-based topology. There are also morphological differences between *I. aguerana* sp. nov *and I. impressa* sp. nov. with the dispar-group sensu Schönhofer et al. (2015). Particularly with the northern Spanish taxa studied by Luque & Labrada (2012), which allow these taxa to be distinguished based on morphology, including the shape of the glans penis. For example, on the dorsal side, the glans has a mass of bristles at its end, arranged into two separate vertical rows that have a particular number and position for each species (Figs. 2, 6, 9): *I. navarrensis* bear only 17 *setae* each side, *I. impressa* sp. nov. 21 *setae*, *I. cantabrica* 25 *setae*, *I. magdalenae* 29 *setae, I. dispar* 31 *setae* and *I. aguerana* sp. nov. 35 *setae*.

Within the Iberian-Cantabrian clade, two main morphological groups can be recognized: the *dispar*-group and the *nodifera*-group. The *dispar*-group includes *I. dispar*, *I. magdalenae*, *I. navarrensis*, *I. cantabrica*, *I. aguerana* sp. nov. and *I. impressa* sp. nov. while the *nodifera*-group includes *I. nodifera*, *I. hispanica*, *I. petiginosa* and *I. damiani* sp. nov. *Ischyropsalis impressa* sp. nov. and *I. aguerana* sp. nov. share diagnostic characters, such as the morphology of the chelicerae and the glans, with the dispar-group as defined by Schönhofer et al. (2015). In these species, the chelicerae apophysis is large, triangular, and pointed with a rounded edge in lateral view, and the male penis glans bears a dorsal mass of bristles arranged into two distinct vertical rows, with species-specific numbers of setae.

By contrast, *I. damiani* sp. nov., is highly similar to other *nodifera*-group taxa. the cheliceral apophysis and penis glans morphology are highly similar across species in this group, as previously noted by Schönhofer *et al*. (2015) making species delimitation based solely on morphology difficult without supporting molecular data. This morphological overlap led Martens (1969b) to synonymize *I. hispanica* and *I. petiginosa* with *I. nodifera*, while Prieto (1990a) later reinstated I*. hispanica* as a valid species, treating it as distinct from *I. nodifera*.

In addition to this, *I. damiani* sp. nov. also present morphological similarities with this group of species, especially regarding glans morphology. All these taxa belong to the Iberian-Cantabrian subclade, whose internal clades are rarely species-specific when only a single marker is used to build the topology, a pattern also reported by Schönhofer et al. (2015) in their first phylogeny of *Ischyropsalis*. The species-specific clades recovered in the combined COI×EF1-α dataset correspond to *I. petiginosa* and *I. damiani* sp. nov. Interestingly, all four of them have a close phylogenetic relationship, especially in the case of *I. petiginosa* and *I. damiani* sp. nov. However, the fact that these two taxa present two different life-styles as *I. petiginosa* has an epigean behaviour (Luque 1991; Luque & Labrada 2012), further supports our view that both are valid species.

A distinct clade is also recovered for *I. hispanica* together with another one for *I. nodifera*, although one *I. hispanica* specimen clusters within the latter in the COI×EF1-α dataset. In the EF1-α phylogeny alone, *I. nodifera* does not form a species-specific clade, while *I. hispanica* does. Further analyses are needed to better resolve the relationships among these two non-troglobitic species. The systematic status of *I. nodifera* and *I. hispanica* remains contentious due to overlapping morphological characters and conflicting interpretations.

SEM analyses (Schönhofer *et al*. 2015) revealed striking similarities in chelicerae microstructure and bristle field, supporting Martens’ synonymy hypothesis (Martens, 1969b) and underscoring the morphological cohesion of the *nodifera*-group. In this context, a detailed integrative reassessment of other nominal taxa, such as *I. asturica*, including topotypic sampling and molecular data, would be highly desirable to clarify their taxonomic status and to resolve long-standing uncertainties within the group.

The existence of two Iberian subclades (The Iberian Pyrenean and the Iberian Cantabrian ones), suggests two different biogeographic lineages: one occurring in the mountainous systems of the Eastern Iberian Peninsula (Schönhofer *et al*. 2015) and another whose members are present in the mountainous systems of the Western Iberian Peninsula (Luque & Labrada 2012). These two lineages would be sister groups and would have a close phylogenetic relationship with the Alpine clade, thus suggesting that a similar speciation process could have taken place in these three lineages.

Climatic oscillations during the Pleistocene, particularly the glacial–interglacial cycles, are known to have played a major role in shaping patterns of diversification in arachnids due to their restricted mobility (Pinto-Da-Rocha *et al*. 2007). Glacial advances and retreats repeatedly fragmented and isolated populations, promoting allopatric speciation across a wide range of habitats. Although these processes affected many epigean species, their impact was especially pronounced in hypogean taxa, where caves acted as stable refugia amid climatic instability (Hedin 1997; Culver & Pipan 2009; Ribera *et al*. 2010; Schönhofer & Martens 2010). Similarly to the case of the southern alpine taxa, the speciation process in northern Cantabrian Range is generally considered to be due to glacial cycles during the Quaternary period, which caused repeated isolations in non-glaciated ‘karst refuge areas’ (Schönhofer & Martens 2010; Luque & Labrada 2012). The preference of *Ischyropsalis* species for permanently cool and humid environments, would have favoured the colonisation of caves (Mammola *et al*. 2019) as their fairly stable climate conditions would have meant that these were the only refuges during the cold periods of the Quaternary. The karst caves inhabited by *I. gigantea* are a clear example as their average temperatures were recorded within the range of 6-9 °C in the Recuistro Cave up in the Miera mountains at 1.300 m a.s.l. (C.G. Luque, unpublished observations). *Ischyropsalis cantabrica*, *I. gigantea* and our three new species are the westernmost cave-dwelling representatives of Ischyropsalidinae, which may have become isolated due to their allopatric ranges (Fig. 1), as the closest karst areas to the localities of these three new species lie on the Sierra Salvada Massif (Burgos–Álava), inhabited by *I. dispar* and Triano-Galdames Mountains (Biscay), inhabited by *I. magdalenae* (Fig. 17). In this context, the extensive glaciation in the Cantabrian Mountains Range was controlled not only by the preexisting landforms and the altitude, but also by the available humidity, related to their distance from the sea (Santos *et al*. 2013; Serrano *et al*. 2013, 2017). This led to species isolation and small-scale endemism and might have been an important factor promoting divergence on karst areas in northern Spain (Sendra 2023). The species delineation presented here provides additional support for this hypothesis and suggests the need for further research on populations across the broad and fragmented distribution range of the genus in northern Spain (Luque & Labrada 2012).

**Figure 17.**
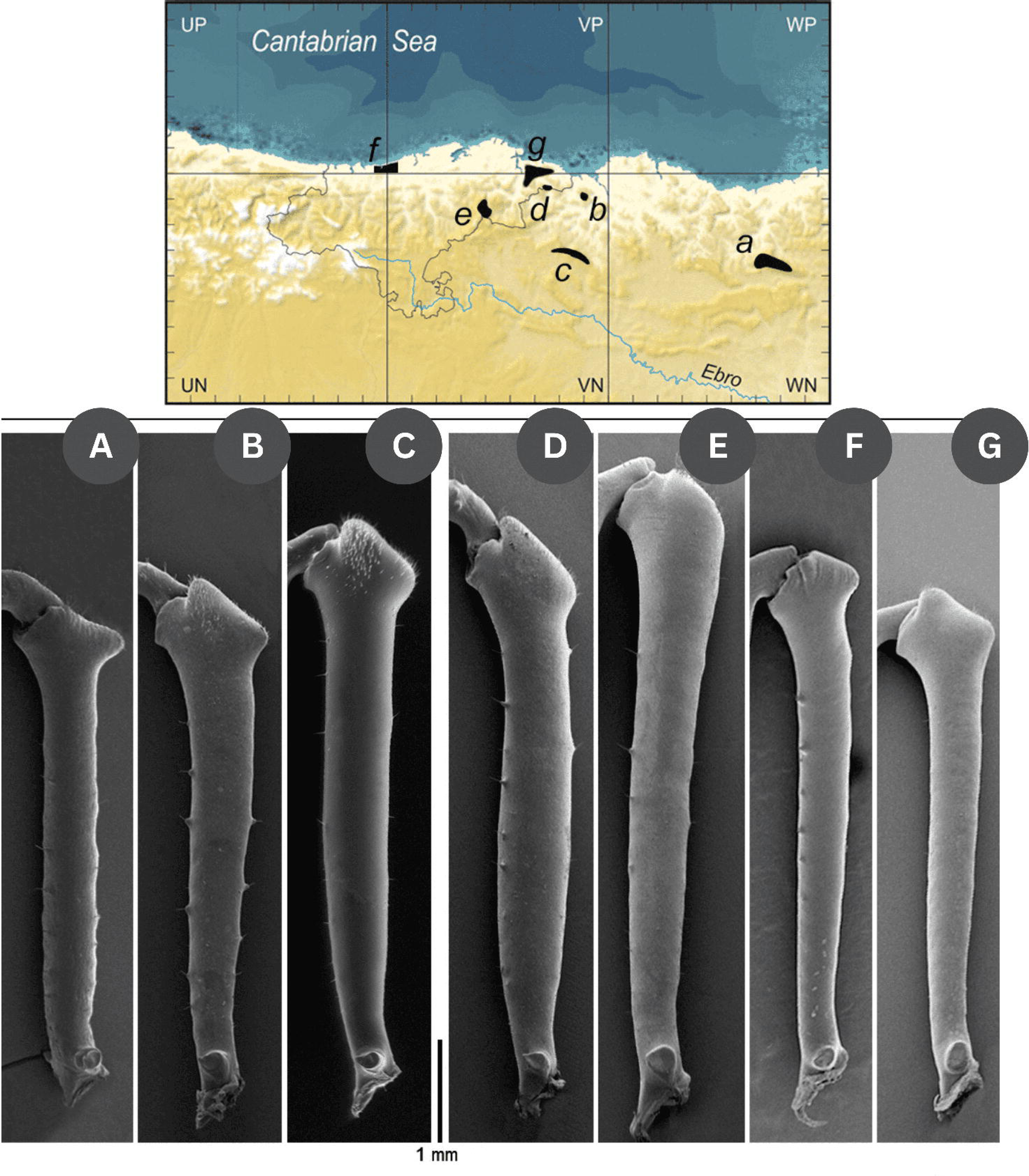
Distribution of the seven Cantabrian troglobiont *Ischyropsalis* species in the Cantabrian Mountains, northern Spain and basal cheliceral article of male (medial view): a. *I. navarrensis*, b. *I. magdalenae*, c. *I. dispar*, d. *I. aguerana* sp. nov., e. *I. gigantea*, f. *I. cantabrica*, g. *I. impressa* sp. nov.; for details, see Luque and Labrada (2012).

In Cantabria and Asturias, the most common conservation strategy for the subterranean ecosystems is focused on the establishment of protected areas (Adrados-González 2011; Ballesteros *et al*. 2013; Luque & Labrada 2015, 2022). Regarding troglobiont *Ischyropsalis* species, only two such species are found in protected areas designated under EU Natura 2000 network (declared December 2004): *I. cantabrica* that inhabits the Rogeria Cave SAC, and *I. gigantea* present in the Eastern Mountain SAC, a group of mountains in the Upper Asón and Miera Rivers Basins. Here are located the largest and most important caverns in Spain and they harbour a wide diversity of poorly understood specialized organisms that are of interest from both a conservational and evolutionary perspective (Labrada *et al*. 2010, in press; Salgado *et al*. 2012; Angus *et al*. 2012; Luque & Labrada 2012, 2017; Faille *et al*. 2021; Fresneda *et al*. 2025).

## Acknowledgements

A posthumous gratefulness to dear departed friends: María Rambla Castells (1917-2016), a professor at the University of Barcelona, for the opportunity to study her interesting materials, which served as the base for the present paper; Miguel Villena Sánchez-Valero (1961-2008), a curator at the National Museum of Natural Sciences (Spain), who kindly provided us with the voucher numbers for our first samples deposited in the museum collection. We are especially indebted to José Bedoya Romero «Josefo» (1962-2002), technician at the National Museum of Natural Sciences (Spain), who took the beautiful SEM micrographs for this work and passed away in a traffic accident in Madrid. We dedicate the present work to the memory of them all. We thank Begoña Sánchez Chillón, a curator at the National Museum of Natural Sciences (Spain), Helena Basas Satorras, a curator at the Animal Biodiversity Resource Centre at Barcelona University, and María Araceli Anadón Álvarez, former curator of the Zoology collection of the University of Oviedo, for enabling the access to the collections, the loan of specimens and providing us with the reference numbers of the collections. We particularly thank to Enric Peradalta Rocha, Administrator of the Speleo-Imserso Project, for providing the image of the Covacho del Cogorio Spring Cave. We also particularly thank to Emilio Muñoz Fernández, Administrator of the project to assess the current state of Palaeolithic rock art conservation in Cantabria’s non-tourist caves, for providing very interesting data and collecting specimens from these caves, we can understand better current patterns in species distributions. We are also indebted to the former members of the Cantabria’s Archaeological Association (CAEAP, Spanish acronym) and SEII-Caving Club (from the School of Industrial Engineering at Madrid’s Polytechnic University) for allowing Carlos G. Luque to accompany them in and around Hoyo Molino, Hoyo Menor and Hoyo de las Fuentes in the 1980s. We want to acknowledge Ana María Arango González for her English revision and correction of the manuscript. Finally, we appreciate helpful comments from the editor and two reviewers.

## Data availability

The data used to support the findings of this study are included within the manuscript. The DNA sequences generated for this research have been uploaded to GenBank (https://www.ncbi.nlm.nih.gov/genbank/).

## Funding

This study was financed by Department Forestry and Nature Conservation of the Regional Government of Cantabria (Contract No. 2004.1.05.03.0025). The cost of technical assistance will be met from heading 05.06.533A.640.02 of the expenditure budget of the regional government (see page 7525 of the Official Gazette No. 141, dated 20 July 2004, www.boc.cantabria.es). Also funded by the previous administration, contract No. 2006.1.05.03.0028. A project cost will be met from heading 05.06.456C.616 of the expenditure budget of the regional government (see page 2071 of the Annual Account Report for the 2006 fiscal year, Vol. 6, Contract Awards, available from https://www.cantabria.es/web/intervencion-general/cuenta-general, last consult Sep. 22, 2021). The funder had no role in study design, data collection and analysis, decision to publish, or preparation of the manuscript.

## Competing interests

The authors have declared that no competing interests exist.

## Author Contributions

Ricardo López-Alonso (conceptualization, methodology, formal analysis, investigation, writing—original draft preparation, writing—review andediting), Esteban Pascual-Parra (conceptualization, methodology, formal analysis, investigation, writing—original draft preparation, writing—review and editing), Lucia Labrada (conceptualization, methodology, writing—original draft preparation, writing—review and editing), Carlos G Luque (conceptualization, methodology, writing—original draft preparation, writing—review and editing), Andrés Arias (conceptualization, writing—review and editing, funding acquisition), Eduardo Cires (conceptualization, writing—review andediting, supervision, funding acquisition), and Claudia González-Toral (conceptualization, methodology, formal analysis, investigation, writing—original draft preparation, writing—review and editing).

## Notes

### Competing Interest Statement

The authors have declared no competing interest.

